# NMRFx: Integrated Software for NMR Data Processing, Visualization, Analysis and Structure Calculation

**DOI:** 10.1101/2025.08.26.672401

**Authors:** Ellen Koag, Simon G. Hulse, Gregory L. Helms, Kevin M. Call, Michael F. Summers, Jan Marchant, Bruce A. Johnson

## Abstract

NMR spectroscopy is applied across a wide range of scientific disciplines to derive chemical, structural, and dynamical information for a broad and diverse range of molecular systems. The utility of the technique depends on robust computational protocols for processing, visualizing, and analyzing a wide range of experimental data types and transforming the data into useful chemical and structural information. Here we introduce NMRFx, a novel software application that integrates and augments features of our existing NMRViewJ and NMRFx Processor applications. NMRFx enables data processing, peak picking and assignment, chemical shift and molecular structure calculation, and beyond, through a high-speed, feature-rich graphical user interface. This paper describes advances over existing software and presents a series of case studies that demonstrate its utility in diverse contexts. These case studies include the assignments of the protein ubiquitin, a 36 nucleotide RNA construct, and the natural product taccalonolide E; and a metabolomics study of triacylglyceride production in algal cells.

## 1 Introduction

Nuclear magnetic resonance (NMR) is a powerful analytical technique with applications in various disciplines including organic chemistry [1, 2], inorganic chemistry [3], environmental chemistry [4], materials science [5], and medical imaging [6]. Since the first determination of a solution-state protein structure in 1985 [7], NMR spectroscopy has been widely used in the field of structural biology [8]. The scope of NMR applications for macromolecular structure analysis continues to grow, with notable advances recently made in RNA research [9], the study of intrinsically disordered proteins [10], and the emerging field of integrated structural biology [11, 12, 13]. Structural biologists also employ NMR to investigate ligand binding [14], metabolomics [15, 16], and molecular dynamics, for which established protocols probe motions over picosecond to multisecond time scales [17].

Central to these applications is the need for robust computational tools for data processing, visualization, and analysis. A widely used software application for the visualization and analysis of macromolecular NMR data is NMRView [18] along with its Java-based successor, NMRViewJ [19]. These highly cited programs have been instrumental in myriad investigations. A complementary program, NMRFx Processor [20], was developed more recently to exploit recent computational hardware advances and provide a feature-rich yet accessible means of processing NMR data.

Here we introduce NMRFx, a program that merges and greatly extends the capabilities of NMRFx Processor and NMRViewJ. Additionally, NMRFx provides tools for structure calculation and allows the inclusion of plugins to enable additional functionality, with a prime example being RING NMR Dynamics [21], an application for the analysis of macromolecular dynamics by NMR. The suite of tools available in NMRFx, which facilitates a complete workflow from the free induction decay (FID) to processed and analyzed spectra, ultimately provides structural and dynamical insights. Although there are alternative programs for NMR processing [22], visualization and analysis [23, 24, 25], and structure calculation [26, 27], none integrate such a wide range of features. Consequently, NMRFx is also appropriate for use in disciplines beyond structural biology. The software is designed to be user-friendly and appropriate for educational settings, allowing users to visualize and adjust various data processing routines in real time.

A summary of the primary features of NMRFx is presented in Section 2. Additional features for specialist use cases, such as support for ligand and pressure titrations [28] and ZZ-exchange spectroscopy [29], are described at https://nmrfx.org. Subsequently, Section 3 provides four case studies that highlight the utility of the software in diverse contexts:

1. Assignment, secondary structure prediction, and dynamics analysis of the backbone atoms in ubiquitin.
2. Assignment of the natural product taccalonolide E.
3. Monitoring carbon fluxes in triglyceride synthesis in an alga.
4. Assignment and structure calculation of a 36 residue RNA.

## 2 Features

### 2.1 Software Architecture: Languages and Platform Compatibility

NMRFx is an open source and extensible cross-platform application. Written in the Java programming language and bundled with the Java runtime environment, NM-RFx can be installed, executed, and developed on all popular operating systems. The graphical user interface (GUI) is built using the JavaFX toolkit [30]. The software is designed for interaction through the GUI for most use cases. It is also possible to issue commands from the command line, allowing for automation of repetitive tasks and delegation of computationally intensive tasks to remote resources. For example, data processing scripts can be written in the Python language, which are executed using an embedded Jython interpreter [31].

### 2.2 Database and File Formats

A key design choice in developing NMRFx has been to prioritize support for recognized file standards over proprietary ones. This ensures seamless interaction with other programs — while NMRFx is designed as a comprehensive tool for NMR analysis, it is easy to import and export data to and from other applications, enabling specific routines to be performed outside of NMRFx. NMRFx accommodates commonly used data formats for both raw and processed NMR data, including Bruker, Varian, JEOL, and NMRPipe. Support is also provided for various data deposition and retrieval formats, such as NMR-STAR [32], NEF [33], PDB [34], and PDBx/mmCIF [35]. The NMR-STAR format is the primary method for storing NMRFx project data. Users can search for and retrieve BMRB entries within NMRFx, and NMR-STAR files can be uploaded for new depositions directly within the application.

### 2.3 Version Controlled Projects

NMR projects often involve numerous intricate steps to process and analyze data, which can span months or even years. By integrating the Git version control system, NMRFx provides features to manage projects of varying complexity. A comprehensive history of all pertinent information, including processed datasets, peak lists, assignments, and window configurations, is maintained in a memory-efficient manner as a tree of snapshots called “commits”. Users can navigate between commits, and create multiple branches from a given commit, allowing for the exploration of alternative protocols within the same project without the risk of losing previous work. Noteworthy commits, such as a project’s state at the point of deposition or publication, can also be tagged for easy recall.

### 2.4 Plugins

NMRFx facilitates the integration of plugins — comprehensive tools written in Java which can be developed and distributed independently of the main program. A notable example of this is RING NMR Dynamics, an application designed for analyzing various types of macromolecular NMR dynamics data, including CPMG, CEST and *R*_1_*_ρ_* experiments, as well as model free analysis [21]. While RING can function as a standalone application, its integration as a plugin within NMRFx enables communication between the two programs, creating a seamless pathway from raw NMR data to dynamic insights. Plugin users can process, peak pick, and assign spectra with the tools of NMRFx (*vide infra*), and subsequently transfer peak information directly to RING for analysis. Additionally, selecting data entries within RING instructs NMRFx to display the relevant spectral region, as illustrated in Figure 6.

### 2.5 Machine Learning Support

As with virtually all scientific disciplines, it has been demonstrated that machine learning methodologies can aid the field of NMR in numerous ways[36].

To facilitate the use of trained models, the TensorFlow library — one of the most widely used deep learning frameworks — is embedded within NMRFx [37, 38]. Furthermore, NMRFx also ships with the Tribuo library, which supports a wide range of classical machine learning models [39]. This enables the seamless introduction of novel models into NMRFx; some examples, including secondary structure prediction of both proteins and RNA, which already ship with the application are described below.

### 2.6 NMR Processing

NMRFx offers a diverse array of operations for processing NMR datasets of arbitrary dimension. Commonly employed operations such as apodization, zero-filling, Fourier transformation, and phase correction are supplemented with operations for specialized use cases including baseline correction, peak suppression, and reconstruction of non-uniform datasets. In most cases an appropriate processing routine is automatically generated through analysis of the dataset’s acquisition parameters, allowing the user to focus on fine-tuning the routine. Wherever possible, each operation is parallelized across multiple CPU cores to ensure computational resources are leveraged to their fullest extent. Routines can be modified within the GUI by interacting with the Processor accordion — a series of expandable tabs ordered by data dimension which outlines the operations to be executed. Operations can be adjusted by interacting with GUI elements, which modify associated parameters. For many operations, the effects of their addition/adjustment are depicted instantly, helping users design optimal processing schemes, while also making NMRFx a valuable educational tool.

Figure 1 shows a screenshot of NMRFx after processing a HNCACB dataset which featured non-uniform sampling (NUS). The processing accordion is expanded to show the options for GRINS (GRINS is not SMILE), a NUS reconstruction algorithm developed within the Johnson group, which has similarities to Sparse Multidimensional Iterative Lineshape-Enhanced (SMILE) reconstruction [40], hence its recursive acronym. In-house implementations of iterative soft thresholding (IST) [41] and NESTA [42] are also available within NMRFx for NUS reconstruction. Interacting with the Processor accordion updates a Python script, which is executed when the user clicks Process. The script used to process the HNCACB dataset is provided in the Supplementary Material (Code Listing 1).

**Figure 1:**
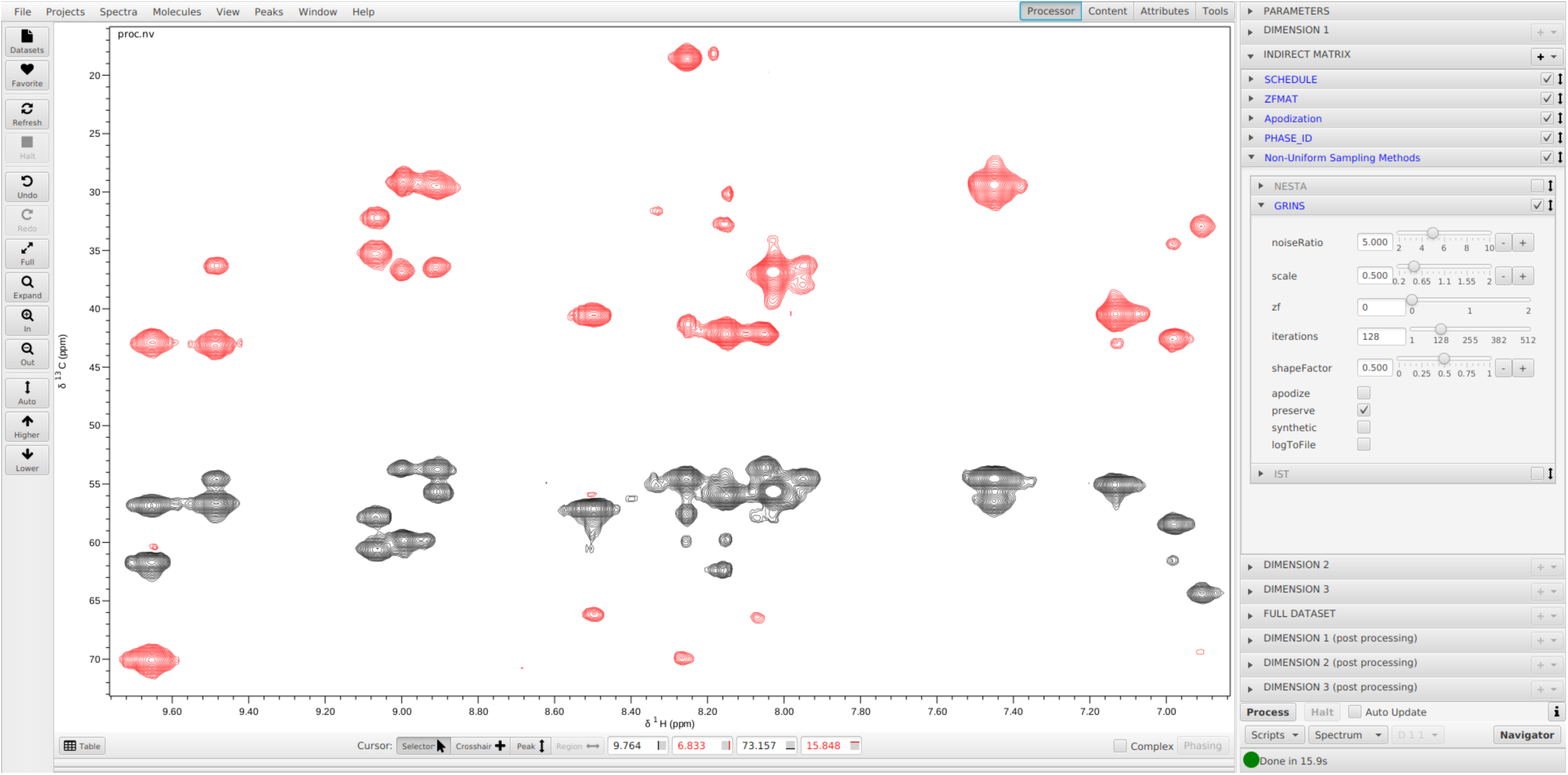
Screenshot of the NMRFx GUI running on an Ubuntu 22.04 system with the GNOME desktop environment. A ^1^H-^13^C plane of an HNCACB spectrum is displayed, with the ^15^N shift set to 119.33 ppm. The spectrum was generated by processing a NUS dataset using the GRINS algorithm. The processing accordion, located on the right-hand side of the GUI, allows users to customize the sequence of steps involved in transforming the raw FID into the final spectrum. Each processing step has an accompanying drop-down menu, with that for the GRINS algorithm visible. The adjustment of processing operations updates a Python script which can be inspected by clicking Scripts > Show Script. The script used to process the presented dataset is listed in the Supplementary Material.

### 2.7 Visualization of Spectra

Correlating data from different NMR experiments is often essential to obtain insights into the chemical system under study [43]. A notable advancement in the original NMRView software was the ability to simultaneously visualize multiple spectra and/or different arrangements of spectrum dimensions in separate windows [18]. This concept has been retained in NMRFx; a virtually unlimited number of top-level windows can be created, with each window able to support numerous spectrum charts arranged within a grid. Correlated cross-hair cursors facilitate the identification of common spectral features.

This capability, while powerful, can be time-consuming to configure. To streamline the process, NMRFx now supports layout files written in YAML, which enable window layouts to be loaded automatically. Users can create custom layouts for their specific requirements. Figure 10 provides an example in which a chart layout has been employed to view correlations between ^1^H-^1^H NOESY and TOCSY spectra, as well as a ^1^H-^13^C HMQC spectrum for an RNA construct. The relevant YAML file can be found in the SI (Code Listing 2). The RunAbout assignment tool (*vide infra*) also makes use of the layout feature to display multiple views of complementary spectra simultaneously, as shown in Figure 4.

### 2.8 Peak Picking

NMRFx provides the same suite of tools for locating (picking) spectral peaks as were present in NMRViewJ, with support present for 1D and multi-dimensional spectra, including pseudo three-dimensional (3D) spectra. It is possible to pick peaks automatically using the built-in peak picker, or interactively pick and adjust them through mouse-based control, which can be useful when peak overlap makes automated picking challenging. Peak features can be refined using non-linear regression tools in order to derive accurate quantitative information such as positions, volumes, and widths. There are recent developments in peak picking using both deep learning [44] and deconvolution-based approaches [45]; the development of similar capabilities is ongoing and will be available in a forthcoming release of NMRFx.

### 2.9 Peak Assignments

NMRFx includes various tools to facilitate the process of assigning spectral peaks to specific atomic sites. These tools aim to automate as much of the assignment process as possible, while allowing user interaction as a means of validation, particularly for more challenging cases. Two specific tools which have found considerable use are the peak slider tool [46], and RunAbout [19].

#### 2.9.1 The Peak Slider

The peak slider tool aids in the assignment of spectra in situations where an appropriate initial guess exists as a starting point [46]. The initial guess typically comes from previous assignments of comparable structures or chemical shift predictions (*vide infra*). Use of the slider tool involves the manual adjustment of peak-boxes to ensure agreement with the spectral peaks. When a peak-box is moved, all other peak-boxes that share an assignment, even those present in other spectra, are simultaneously updated, ensuring consistency and reducing ambiguity. Furthermore, when a peak-box is deemed to be correctly positioned, it can be frozen in place. The dimensions of other peak-boxes assigned to the same atom are also frozen, such that they may only be adjusted in the remaining free dimensions. This mechanism is illustrated in Figure 11 and its use described further in Section 3.4.

#### 2.9.2 RunAbout

The assignment of triple resonance spectra of proteins using NMRFx is facilitated by RunAbout, which was introduced as part of NMRViewJ [19]. In NMRFx the tool has been optimized to use the new GUI toolkit and to be more flexible in terms of what experiments it potentially supports, including those involving direct detection of ^13^C and ^15^N. In addition, more functions that automate the assignment process have been incorporated, minimizing the need for manual interaction. The operation of RunAbout consists of three main operations:

1. **Grouping spectral peaks**: Peaks are grouped into distinct “spin systems” — clusters of peaks that share the same root frequencies, typically those of the backbone H^N^ and N spins.
2. **Linking spin systems**: Adjacent spin systems are connected to form contiguous fragments through the consideration of various criteria.
3. **Matching fragments** The fragments are aligned with regions of the protein’s primary sequence to create the assignments.

RunAbout allows users to quickly navigate the sparsely populated 3D spectra, focusing only on areas with significant peak groupings, akin to island-hopping in the boat which is program’s namesake.

NMRFx’s support for complex spectral layouts (Section 2.7) is exploited in Run-About to ensure all spectra can be considered in a holistic manner. When analyzing peaks and grouping them into spin systems, two orthogonal views — ^1^H-^13^C and ^15^N-^13^C — are considered for the intra-residue and inter-residue spectra. These views aid the inspection and fine-tuning of the peak picking.

When linking spin systems, three distinct groupings of spectra are typically presented, as illustrated in Figure 4. The two central columns display the peaks associated with the spin system currently under consideration (*i*), while the two leftmost and right-most columns show the corresponding regions for spin systems which are candidates for *i −* 1 and *i* + 1, respectively. Spectra containing signals for different carbon types (C^O^, C^α^, C^β^) are organized into separate rows. Several metrics are provided to help users decide whether the current candidates for *i −* 1 and *i* + 1 are valid. These include a score quantifying the peak correlations [47], a tally of the number of peaks which are correlated, and boolean flags indicating:

- Reciprocal match: spin system *i −* 1 does not form a better match with another candidate.
- Available for matching: the spin systems considered have not yet been assigned to a fragment.
- Viable fragment: the chemical shifts are consistent with at least one contiguous amino acid pair in the primary sequence.

Additionally, the program presents the chemical shifts associated with the *i* spin system and the *i −* 1 candidate, as well as lists, ordered by likelihood, of possible amino-acid types for the spin systems based on said shifts. Based on all this information, the user can manually confirm the linkages. Alternatively, an automated protocol can use the same criteria to form links across a series of spin systems. Linked spin systems, especially when they form fragments of three or more systems, can then be matched to specific residue positions.

NMRViewJ introduced an automated protocol that combines the formation of spin system links and the matching of the linked spin systems to sequence positions into a single automated process[48]. This process uses a bipartite matching algorithm to generate a population of possible mappings of the spin systems to primary sequence regions. These sets of matches form an initial population with higher matching scores than would be present with a simple randomly generated population. The population of matches are then optimized by a genetic algorithm. NMRFx has an optimized version of this code with improvements in the protocol used to generate the population of matches and the genetic algorithm used, and the process is integrated into the GUI. Although there are many other programs available for automated assignments, including, for example, BARASA [49], FLYA [50], and MARS [51], the close integration of this automation into the NMRFx RunAbout GUI allows the user to use all RunAbout features to inspect, modify, and extend the results of the automated analysis.

### 2.10 Chemical Shift Prediction

The assignment of spectral peaks to atomic sites can be complex and time-consuming. As a result, there has been significant interest in predicting chemical shifts based on molecular structure [52, 53, 54, 55]. These predictions can be used as *a priori* estimates, which can be refined through the consideration of spectral data (recall the peak slider, Section 2.9.1). NMRFx provides a suite of models designed to predict chemical shifts in proteins, RNA, and small molecules. Beyond the descriptions provided here, further information about the models can be found in the Methods section, as well as the Supplementary Material.

#### 2.10.1 Proteins

NMRFx uses a trained linear regression model for protein chemical shift prediction. In a similar fashion to other programs [52, 53, 54], the model makes use of numerous features which are derived from primary, secondary, and tertiary structural information, including dihedral angles, residue attributes, ring current shifts, and hydrogen bonds. Individual models are trained for each pairing of amino acid and atom type (C^α^, C^β^, ^15^N, H^N^, H^α^, etc.) to account for the varying influence of features in each case.

The high dimensionality of the supplied feature vector risks overfitting the training dataset, which would render the model ineffective at general predictions. To account for this, the model is regularized using least absolute shrinkage and selection operator (LASSO), and trained with the least angle regression (LARS) algorithm [56]. Figure 2 compares the performance of the model with the widely used SHIFTX+ program [52], on a test dataset comprising the 61 proteins outlined in Table S2 of [52]. It can be seen that the overall performance of NMRFx’s model is similar to SHIFTX+ across the backbone atom types. Chemical shifts of all C, N and H atoms with sufficient training data can be predicted.

**Figure 2:**
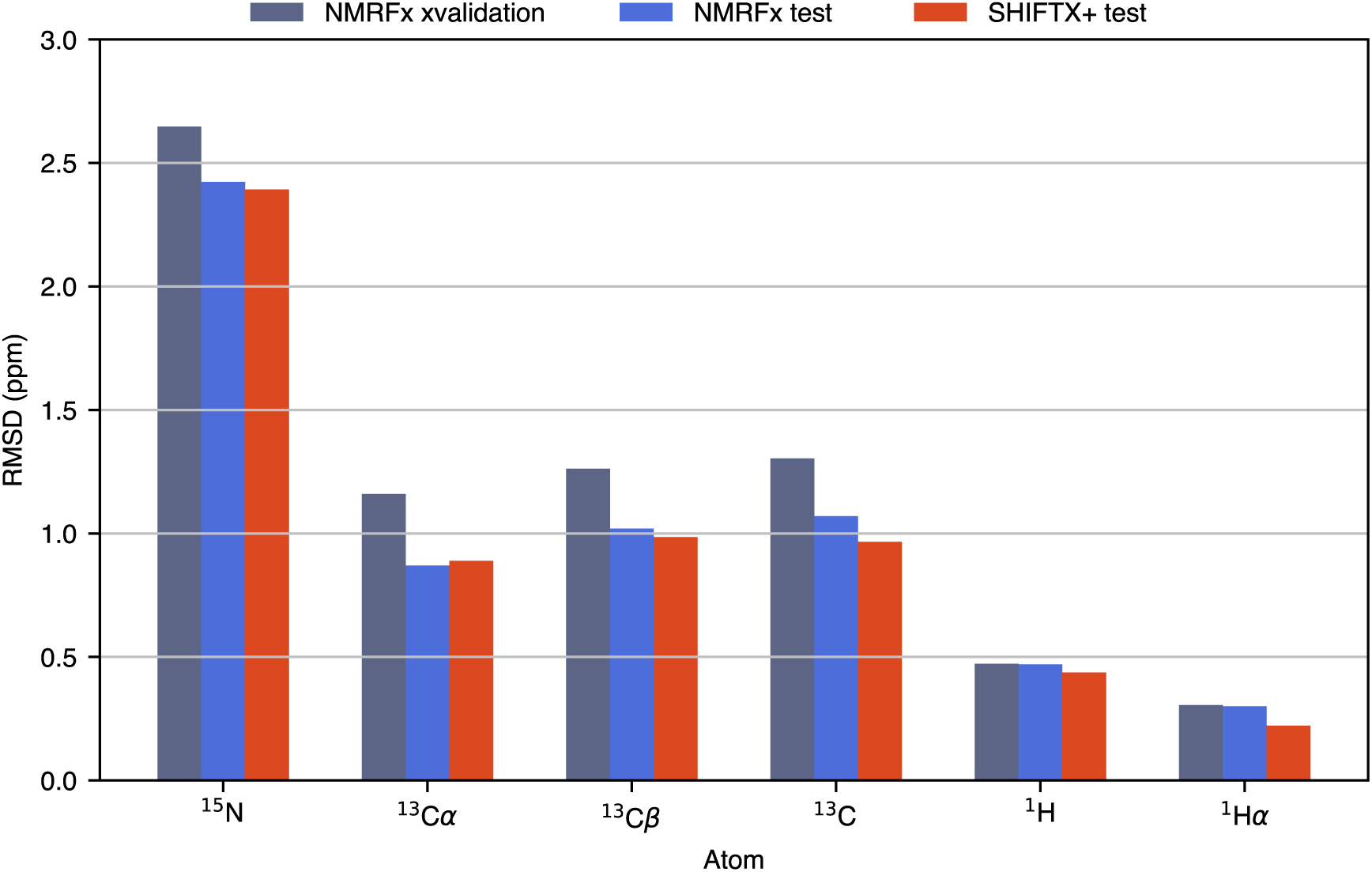
Protein chemical shift performance of NMRFx compared with that of SHIFTX+, a widely-used structure-based shift prediction algorithm. The RMS deviation between the predicted values and experimentally determined values are shown for backbone atoms. The test set used is the 61 proteins originally used for SHIFTX+. The RMSD values for SHIFTX+ are taken from the original publication [52]. Also shown for NMRFx are the values for 10-fold cross validation of the training set.

#### 2.10.2 RNA

NMRViewJ featured RNA shift predictors based on support vector regression (SVR) [57, 58]. NMRFx features new models based on Factorization Machines (FM) [59]. An overview of the prediction performance of the FM compared to the SVR implementation is provided Table 4 of the SI. Features supplied to the model are derived from the primary and secondary structure of the RNA, with considerations made for the base of interest, the two previous and successive bases in the primary structure, and any pairing partners associated with these bases. NMRFX can also predict RNA shifts using 3D RNA structures, employing models that utilize either ring-current shifts [58] or weighted distances of atoms to the target atom, similar to the method introduced by Frank et al. [60].

#### 2.10.3 Small Molecules

Small-molecule chemical shifts can currently be predicted by generating features akin to hierarchical ordered description of the substructure environment (HOSE) codes [61], in which each atom is represented by up to six “shells” of bonded atoms. A database of assigned shifts is then searched, and the average shift of sites with matching representations is provided as the prediction. In a future release of NMRFx, deep learning models are also expected to be available.

### 2.11 Structure Prediction

#### 2.11.1 Structures from NOESY Distance Constraints

NMRFx provides tools for generating 3D molecular structures that are consistent with experimental NMR derived distance and angle restraints. These tools have evolved from PEGASUS [62] which calculates structures in torsion-angle space — a methodology also adopted by CYANA [26] and XPLOR-NIH [27]. NMRFx’s support for a wide variety of file formats affords the user flexibility in incorporating the constraints. Among these, NEF files are recommended for their superior portability with other programs. For example, initial structure predictions can be generated in NMRFx, followed by refinement using programs such as AMBER [63].

Typically, a structure calculation involves generating an ensemble of candidate structures, each optimized to adhere closely to the NMR constraints with low values of the force-field energy. Starting the structure calculations from the provided **nmrfxs** command line application allows the calculation of each structure to run in parallel in a separate operating system process (a feature not yet supported in the GUI). The various parameters for the calculation, such as the number of steps, temperatures, and forces, can be provided in the YAML format.

The structure generator was tested by calculating the 3D structures of proteins in the 100-protein NMR spectra dataset [64]. For each dataset entry, a NEF file containing NMR constraint information was provided as input, resulting in the generation of an ensemble of 200 distinct structures. A cost function was computed for each structure in the ensemble. This function quantifies the degree to which distance and angle constraints were violated, while also including repulsive terms to penalize atom overlap. The 10 structures with the lowest cost function were output to a PDBx/mmCIF file. These 10-structure ensembles were then averaged and superimposed onto the corresponding experimentally confirmed structures obtained from the PDB. The results generated by NMRFx were compared with published structures by computing the RMS deviations between all backbone atoms in the calculated structures and the PDB structures, with the results presented in Table 1 of SI. Nine examples of the results are shown in Figure 3, demonstrating strong agreement. Furthermore, the minimum and maximum violations of each NMRFx calculation are provided in Table 2 of the SI.

**Figure 3:**
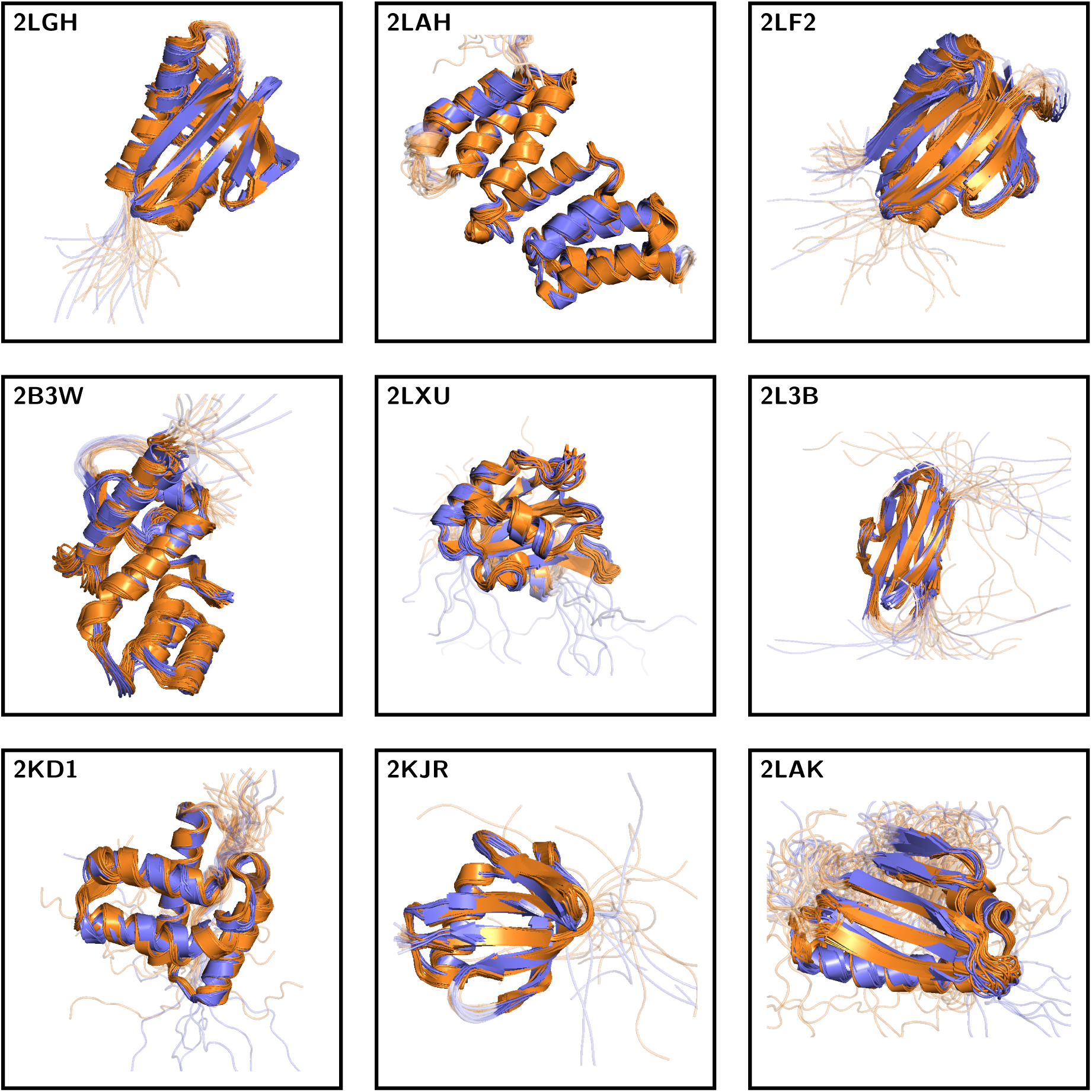
Structure recalculation results generated by NMRFx on a subset of the proteins present in the 100-protein NMR spectra dataset [64]. The 10 best structures sampled from an ensemble of 200 calculations are each shown (purple), and superimposed on the corresponding models (typically 20) from the PDB deposition (orange).

#### 2.11.2 2D to 3D Structure Mappings

NMRFx also provides support to map 2D small molecule structures to 3D structures using the embedded library OpenChemLib [65] which can generate energy optimized 3D structures starting from atom types and bonds in a 2D structure. These structures aid in chemical shift prediction, facilitate the calculation of macromolecular-ligand complex structures, and assist in the assignment of NOESY crosspeaks when used in conjunction with the slider tool.

### 2.12 Sequence Display Tool

NMRFx provides a sequence display tool designed to present residue-specific properties of macromolecules. A variety of parameters can be rendered in bar or dot plot form (Figure 5). These include chemical shift deviations from statistical means or predicted “random coil” values (predicted using an NMRFx implementation of the Potenci algorithm [66]); plots of CheZOD *Z*-scores, which quantify the extent of order versus disorder [67]; and protein secondary structure predictions. The latter predictions are performed with a deep learning model implemented and trained with Tensorflow and integrated in NMRFx. The model is trained to predict a four-state secondary structure prediction based on a subset of states generated by DSSP [68] (see Figure 5 for details).

**Figure 4:**
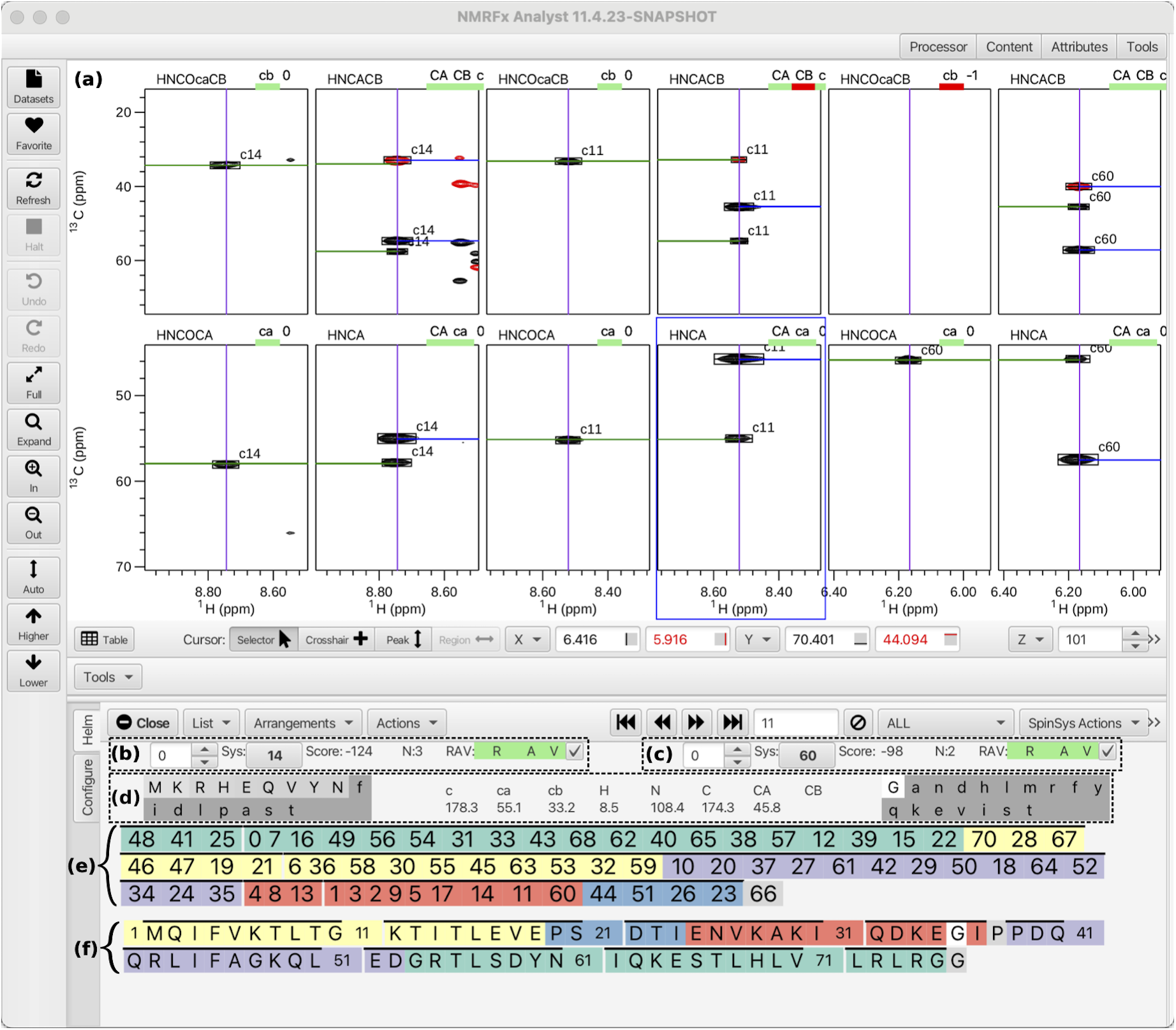
Screenshot of the NMRFx RunAbout tool in the process of assigning ubiquitin chemical shifts, after running the automated assignment algorithm. **(a)** Spectral windows are arranged such that the upper row comprises views of the HNCOCACB and HNCACB spectra, while analogous views of the HNCOCA and HNCA spectra are on the lower row. The central two columns display the peaks which are associated with the current spin-system under consideration (11), while the two left- and right-most columns display those of candidates for the *i −* 1 and *i* + 1 residues, respectively (14 and 60). **(b), (c)** Information related to the candidates. From left to right: a spinner to iterate through the candidates in order of likelihood; the identity of the spin system (Sys); a metric which indicates the goodness of the peak correlations with the *i* residue (Score) [49]; a tally of the number of peaks which are correlated (N); and indicators of reciprocity, availability, and viability (RAV). **(d)** The chemical shifts associated with residues *i* (H^N^, N, C, C^α^, C^β^), and *i −* 1 (C, C^α^, C^β^). Lists of the most likely amino acid types for the *i* and *i −* 1 residues are provided on the right- and left-hand sides respectively; the brighter the shading, the more likely the amino acid. **(e)** Groupings of spin systems that have been sequentially connected, ordered by group length. **(f)** Mappings of the spin system groups onto the protein primary structure, generated using the bipartite matching/genetric algorithm based automated assignment tool.

**Figure 5:**
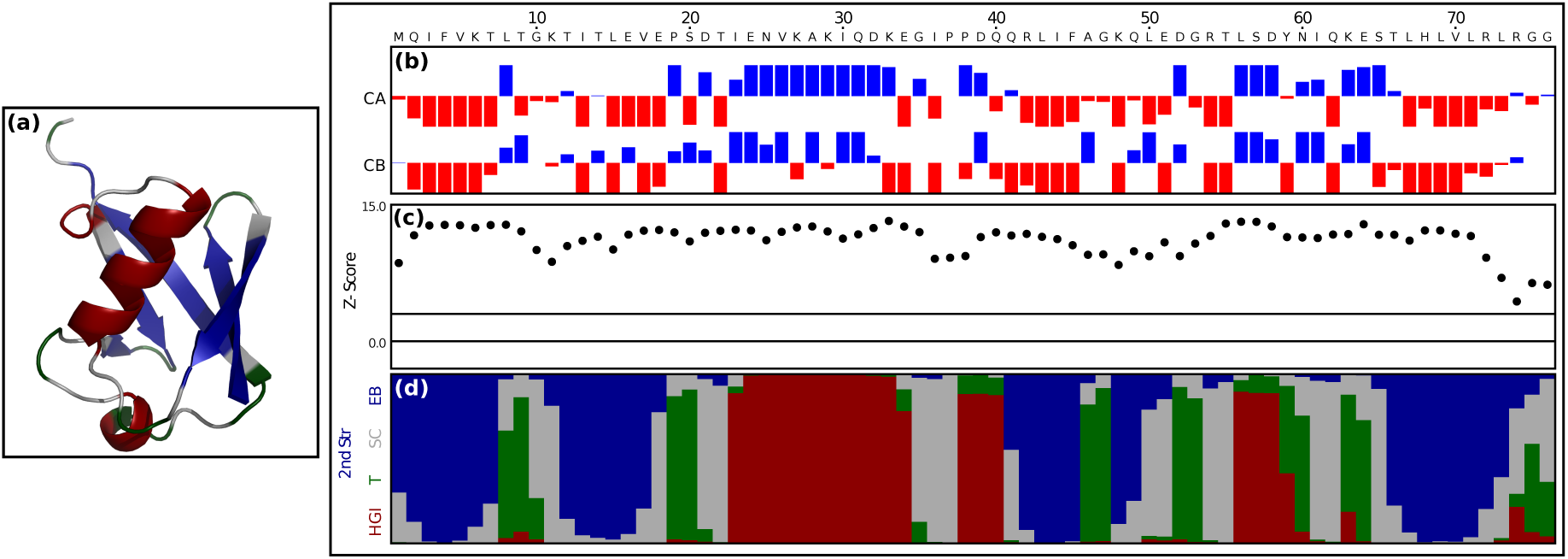
The NMRFx sequence display tool presenting a secondary structure analysis of ubiquitin. **(a)** Cartoon diagram of crystal structure of ubiquitin, refined at 1.8 Å resolution (PDB deposition 1UBQ) [76] **(b)** Deviations of C^α^ and C^β^ chemical shifts from estimated random coil chemical shifts [66]. **(c)** Plot of the CheZOD *Z*-score, an indicator of the degree of residue-specific disorder [67]. As a rule of thumb, residues for which *Z <* 3 are considered disordered (*Z* = 3 is plotted in solid black); those for which 3 *< Z <* 8 are partially formed, and those for which *Z >* 8 are fully-formed. **(d)** Probabilities of residue classifications, using the folowing four-state system: red: HGI (alpha helix); green: T (hydrogen-bonded turn); gray: SC (bend/coil); blue: EB (extended sheet).The carton diagram of panel a is colored such that each residue corresponds to the classification with the highest probability.

## 3 Case Studies

### 3.1 Ubiquitin: Assignment, Secondary Structure Analysis, and Dynamics Analysis

In this case study, a number of NMRFx’s features are employed to perform a detailed study of the backbone atoms of ubiquitin [69].

#### 3.1.1 Backbone Assignment Using RunAbout

RunAbout (Section 2.9.2) was used to perform chemical shift assignments. The assignment was performed twice. In the first case, it was done with automated tools to form spin systems and assign peaks to atom types, followed by manual spin system analysis and confirmation of links between potentially adjacent spin systems. In the second case, linkage and residue assignment was performed with the automated tool (bipartite match / genetic algorithm). The confirmation that the assignments were correct in both cases was made by comparing the assignments with published data (as BMRB files). The user-facilitated process was saved as a Git project, with frequent commits made. This project is available and serves as a useful “follow-along” tutorial for new users of RunAbout. The following summarizes the workflow used.

HNCO, HNCACO, HNCACB, HNCOCACB, HNCA and HNCOCA datasets of ubiquitin were processed and peak-picked within NMRFx. The peak lists were optimized by (a) filtering out peaks that were not consistent across the spectra, (b) clustering the remaining peaks into spin systems, and (c) classifying each peak based on atom type (C^O^, C^α^, C^β^) and connectivity (intra- or inter-residue). In general, particular care was taken when considering the HNCO spectrum, as its generated peak list was used as a reference in order to synchronize with the other spectra.

After the peaks were assembled into spin systems, sequentially adjacent spin systems were identified, making use of the compatibility metrics described above. In this example, chemical shift values,— used to assess likely amino acids for each residue and the correlation between fragments and regions of the primary sequence — were derived from statistical information from the BMRB. Alternatively, they can be derived from known values for a related protein, or predictions based on a known 3D structure using the NMRFx (or other) chemical shift predictor. As spin systems are linked to form longer fragments, the number of viable regions in the primary sequence which are correlated to a given fragment chemical shifts is reduced, until there is typically only a single solution.

Figure 4 depicts the result of the second, automated, analysis. The assignments, with the exception of the final glycine residue, are fully complete and consistent with those reported previously. In general, a single complete fragment for the entire protein is not expected to be formed. Fragments (colored sequences shown in the figure) span regions bounded by proline residues or, in more complex spectra, those with missing spectral information. For more challenging assignment problems, users can make use of a combination of automated tools and manual interactions in difficult regions. This is facilitated by the tight integration of the automated tools into the Runabout GUI.

**Figure 6:**
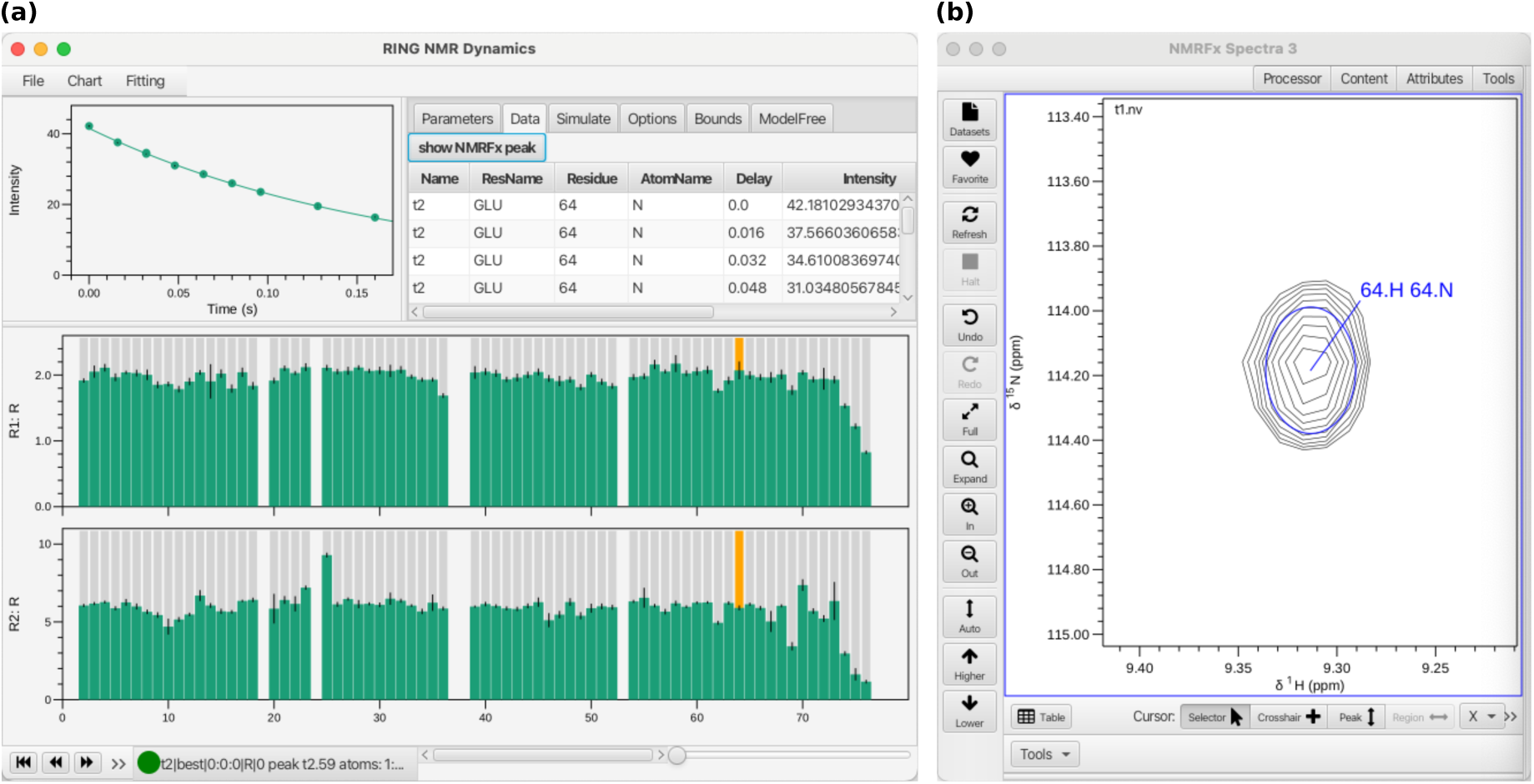
(a) RING NMR Dynamics running as an NMRFx plugin. The bar charts at the bottom of the window show estimated backbone amide ^15^N *R*_1_ and *R*_2_ values of amino acid residues in ubiquitin. Selecting a residue entry (column highlighted in orange) shows a plot of the measured intensities as a function of relaxation delay time in the top left of the window. Selecting an entry in the data table adjacent to the plot, and clicking show NMRFx peak instructs NMRFx to display the relevant peak (panel **(b)**), which in this case corresponds to residue 64 (gluatmic acid) in ubiquitin.

#### 3.1.2 Predicting Secondary Structure

The assigned backbone chemical shifts were subjected to further analysis using the NMRFx sequence display tool (Section 2.12), as shown in Figure 5. Shown are chemical shift deviations to random coil values, an estimate of residue specific disorder, and a four-state prediction of secondary structure.

#### 3.1.3 Dynamics Analysis with RING NMR

RING NMR Dynamics, incorporated into NMRFx as a plugin (Section 2.4), was used to derive backbone dynamic information about ubiquitin, through the calculation of amide ^15^N *R*_1_, *R*_2_, and NOE values. The dataset peaks were assigned to residues using the assignments computed with RunAbout. NMRFx is equipped with a peak simulator, which was used to generate a synthetic ^1^H-^15^N HSQC experiment, using the assigned chemical shifts as peak positions. Subsequently, the associated peak-boxes were interactively aligned with the actual resonance positions in the relaxation datasets, before peak intensities were calculated in each 2D plane of the pseudo-3D datasets. At this point, RING NMR Dynamics was invoked within NMRFx via the Plugin menu, allowing for communication between the two programs. Figure 6 illustrates the result of fitting of the peak intensities to exponential equations, leading to the *R*_1_ and *R*_2_ estimates. Complete analysis of the *R*_1_, *R*_2_ and NOE data can be performed, including model-free analysis to derive order parameters and correlation times.

### 3.2 Assignment of Taccalonolide E

Although NMRFx has been developed predominantly with structural biology applications in mind, its rich set of features makes it a valuable tool for the interpretation, assignment, and structure determination of small molecules such as synthetic compounds or natural products. NMRFx tools for 1D spectra allow facile peak picking, integration, and multiplet analysis in manual and automated modes. Figure 7 shows the result of automated processing and analysis for the ^1^H spectrum of the natural product taccalonolide E [70]. The spectrum is shown in panel a, along with a zoomed-in region in panel b, which includes annotations describing the two multiplet structures present. This analysis is facilitated by simulating the spectrum, followed by fine-tuning the peak features (positions, intensities, widths) with an optimization routine to best match the experimental data. Complex or overlapping multiplets can be individually adjusted by the user so that the components can be accurately described. Furthermore, relevant molecular structures can be viewed alongside the spectral data and assignments made by selecting a multiplet and clicking on the corresponding atom (panel c), and summaries of the spectral data can be generated in a variety of formats supported by different journals (panel d). NMRFx, was used to process and analyze a suite of datasets comprising 1D ^1^H, 1D ^13^C, HSQC, TOCSY, ROESY and HMBC spectra, from which complete peak assignments could be made, including all diastereotopic methylene protons and methyl groups, with the assignments being in agreement with those reported in the literature. It should also be noted that advanced assignment tools such as the slider tool are applicable to small molecules, and their application is presented in the RNA case study (*vide infra*).

**Figure 7:**
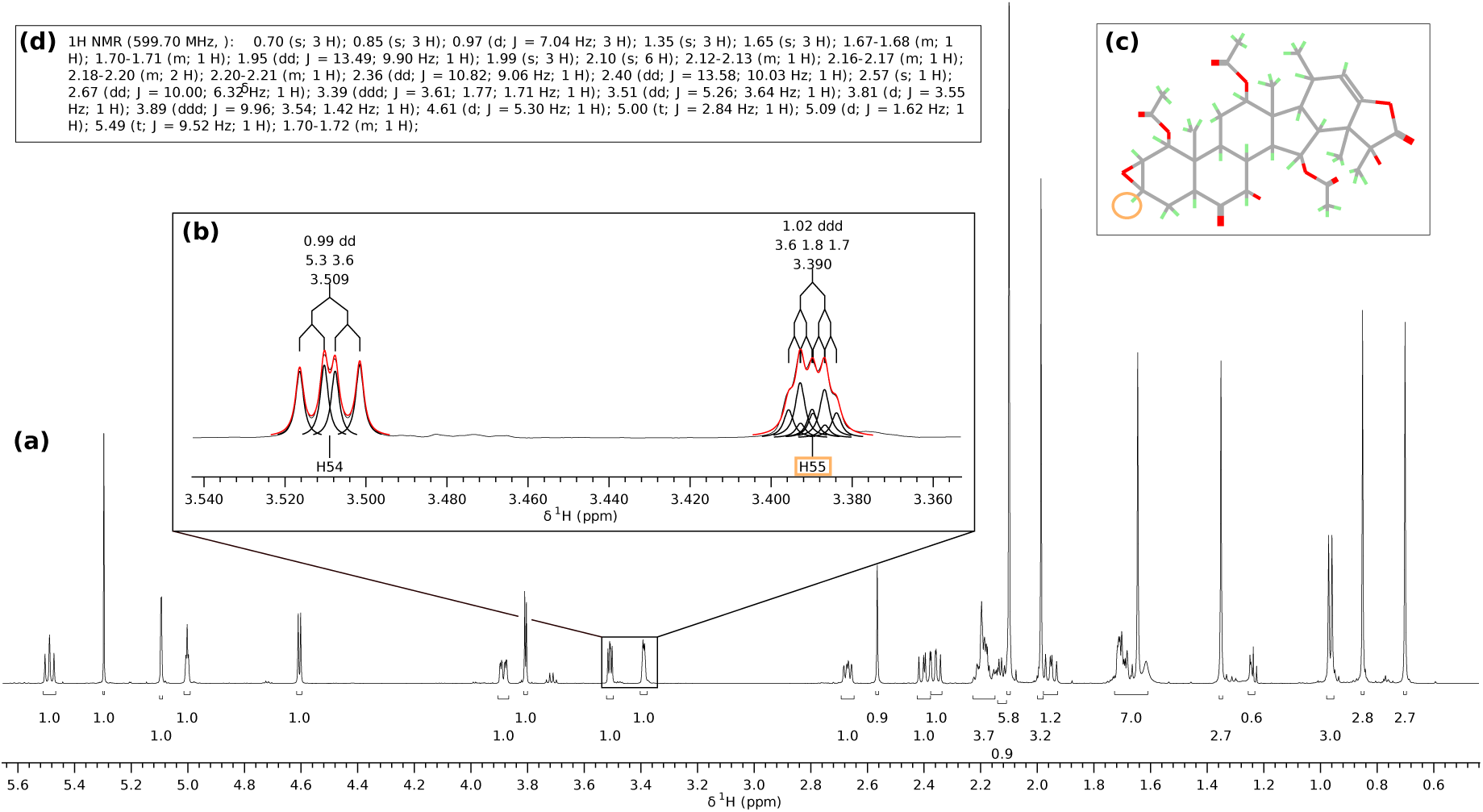
Analysis of a 1D ^1^H spectrum of the natural product taccalonolide E using NMRFx. **(a)** A wide view (5.8 ppm to 0.5 ppm) of the spectrum, with multiplet integrals denoted to 1 decimal place. **(b)** A zoomed-in view (3.54 ppm to 3.36 ppm) of the same spectrum, showing two multiplet structures. The experimental spectrum is plotted (thin black line), along with the result of fitting the multiplets, with individual peak components (thick black lines) and their sum (red lines) shown. Further useful information about each multiplet, including relative integrals (2 decimal places), J-coupling values (Hz), central frequency (ppm), and a tree diagram outlining the couplings, has been displayed. **(c)** Display of the molecular structure of taccalonolide E. It is possible to visualize links between multiplet structures and atomic sites by interacting with the spectrum, with one such link highlighted in orange. **(d)** A generated summary of the spectrum, in J. Org. Chem. format.

### 3.3 Metabolomic Study of Triacylglyceride Production in Live Algal Cells

NMRFx has extensive functionalities for comparing, overlaying and analyzing multiple 1D or 2D datasets, acquired as individual datasets or as arrays, for probing structural analogs, kinetics, relaxation, diffusion, metabolomics, ligand titrations or any spectra collected as a function of one or more variables. The Scanner Tool (accessed via the Tools menu) is versatile for quantifying, visualizing and plotting multiple datasets. Individually acquired datasets or arrayed datasets can be loaded into the Scanner Tool using either the Datasets dialog or via loading a text table (comma or tab separated) with column headers that contain the path to the data, dataset name and any pertinent variables that describe how the data was collected. In the case of metabolomics, data variables may be the time/date of data collection, the replicate number and whether the sample was collected from a control or dosed cohort. Figure 8 shows a metabolomics study of live algal cells where 1D High-Resolution Magic Angle Spinning (HRMAS) ^1^H spectra were acquired by sampling, as a function of time, a culture of *Chlorella vulgaris* deprived of nitrogen and supplied with NaH^13^CO_3_ as the sole carbon source. This study shows the accumulation of triacylglycerides (TAG), a model of which is shown in a), as well as sucrose which increases with time post addition of the labeled bicarbonate. The data were collected in triplicate over 38 hours with broadband ^13^C decoupling (C13 = nny) b) or left ^13^C coupled (C13 = nnn) as in c). The raw FIDs were processed via the Processor and phased and baseline corrected. The Scanner Tool applies the operation to all FIDs in the table. The resulting spectra can then be visualized and grouped in multiple ways as shown with the ^13^C decoupled spectra grouped in red a) and the ^13^C coupled spectra grouped in black b). The spectra are also grouped by replicate and in increasing time of sampling. Spectra can be aligned to account for shifts due to pH changes as well as normalized in a number of ways. Integration across the dataset is accomplished by integrating discreet regions which form new columns in the table. This can be seen as a column in d) labeled Methyl Center vo 0.9457 0.8508 re where the methyl groups of the fatty acid chains are integrated between 0.85 and 0.95 ppm. Each of the distinct resonances of the triacylglyceride and sucrose metabolites were integrated. The resulting integral values stored in the table columns can be plotted and fitted using built-in plotting tools or exported as CSV or TSV files for external analysis. Results for the plot of the saturated CH_2_ resonance integral values for both the decoupled (red) and coupled (black) spectra can be seen in e). The increase in integral area upon decoupling indicates most of the carbon incorporated into the saturated fatty acids is derived from ^13^CO_2_ fixed during photosynthesis. The minimal increase in integral value for the omega-3 methyl group f) upon ^13^C decoupling indicates the majority of the carbon comprising the polyunsaturated fatty acids is derived from cellular carbon pools established before addition of the labeled bicarbonate (carbon recycling). The plot of the integral values for sucrose shows rapid and exponential incorporation of labeled carbon which reaches a steady state after about 16 hours. In addition to the generic plot tool (scatter or box), NMRFx includes plotting tools specific for diffusion data as well as for tracing the trajectory of resonances in 2D ligand titration studies.

**Figure 8:**
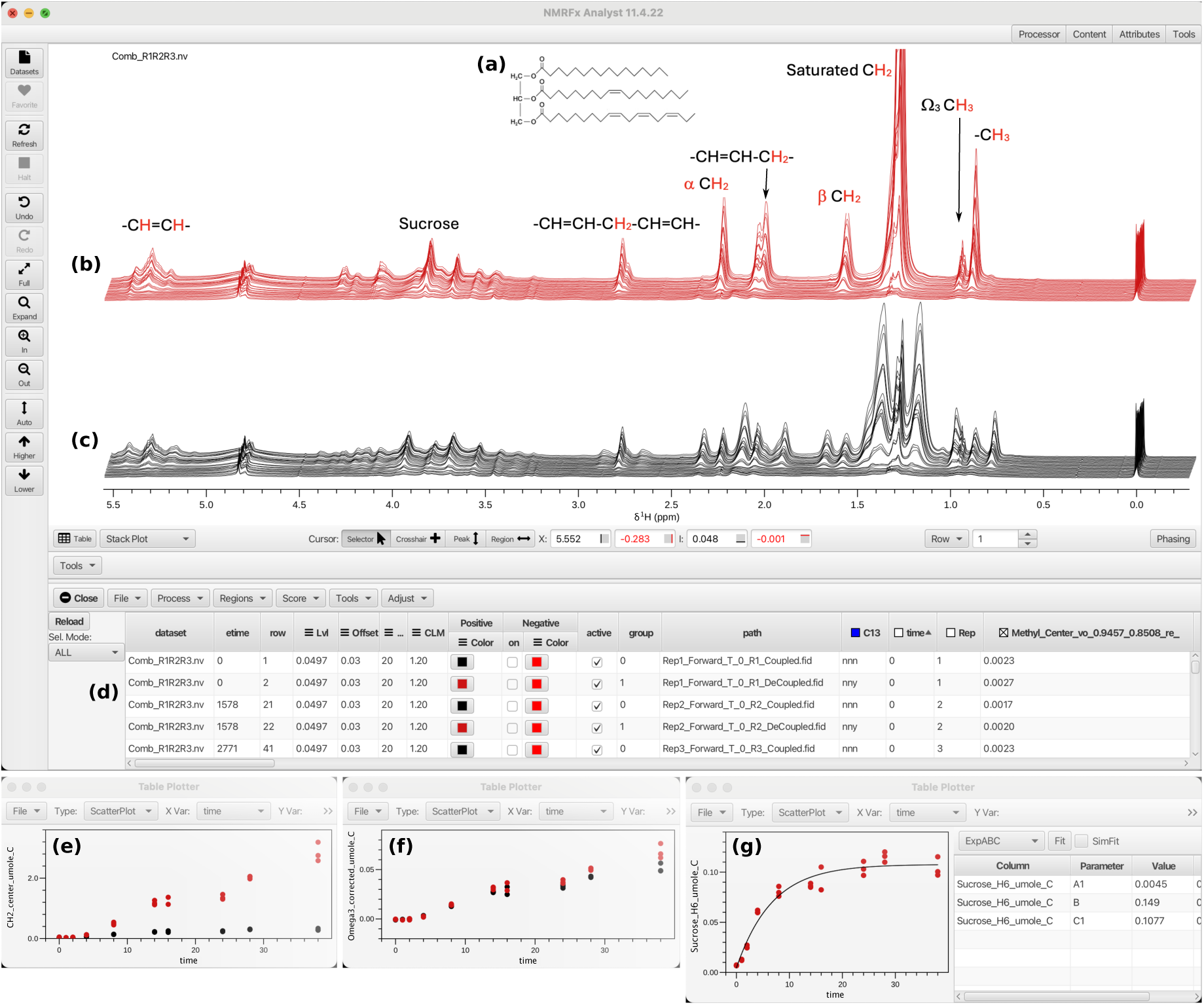
The NMRFx scanner tool displaying a series of 1D ^1^H HRMAS spectra of live algal cells sampled over 38 hours, after being supplied NaH^13^CO_3_ as the sole carbon source. **(a)** Structure of the model triacylglyceride with saturated C16:0, monounsaturated C18:1 and polyunsaturated C18:3 fatty acid chains. **(b)** Broadband ^13^C decoupled ^1^H spectra with fatty acid resonance positions labeled. **(c)** ^13^C coupled ^1^H spectra acquired at the same points over the 38 hour period as the decoupled spectra. **(d)** Control panel for interacting with the scanner tool. **(e)–(g)** Plots of spectrum integral as a function of time for the decoupled and coupled spectra, for the following spectral regions: (e) saturated CH_2_, (f) Ω_3_ CH_3_, (g) Sucrose C6 methylene. The sucrose data in (g) is presented alongside a fit of an inverse exponential of the form *I* = *A* + *B*(1 *− e^−Ct^*), with *A* = 0.0045, *B* = 0.149, *C* = 0.1077 s*^−^*^1^.

### 3.4 Assignment and Structure Calculation of a 36 nt RNA Construct

NMRFx has been used by the Summers laboratory to assign the aromatic and H1*^′^* chemical shifts of more than a dozen RNA constructs with sizes of up to 60 nt. This discussion focuses on a 36 nt RNA construct derived from stem loop C of the MMLV 5*^′^*-Leader (SLC^A^), which consists of two helices separated by a non-canonical k-turn [71, 72]. ^1^H-^1^H NOESY, ^1^H-^1^H TOCSY and ^1^H-^13^C HMQC datasets were processed in NMRFx using built-in scripts with parameters automatically extracted from files within the Bruker experiment directories.

The chemical shift assignment strategy employed here used several NMRFx features, including RNA secondary structure prediction, chemical shift prediction, spectral peak prediction, the ability to link peak-box positions during interactive manipulation (“peak sliding”), and the spectral layout tool.

Chemical shift predictions were generated using NMRFx’s FM model (Section 2.10.2), which incorporates the RNA’s secondary structure of the RNA as input. The secondary structure used in this study was predicted from the primary sequence using a deep learning model implemented within NMRFx with Tensorflow; the secondary structure is presented in Figure 9. Optionally, an explicit secondary structure can be provided in dot-bracket notation [73].

**Figure 9:**
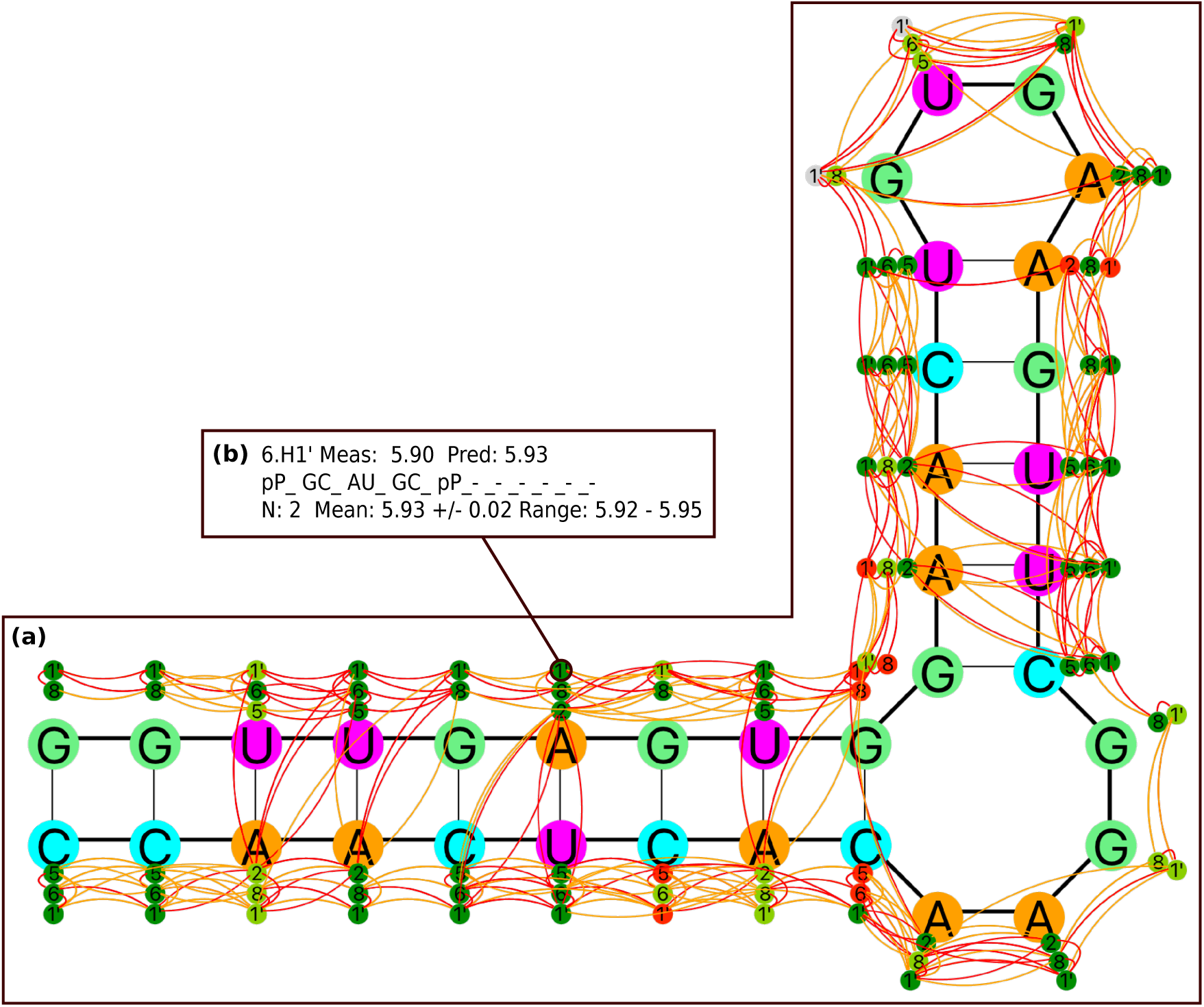
The RNA secondary structure display tool in NMRFx, showing the 36 nt RNA construct considered. **(a)** The predicted secondary structure, generated using NMRFx’s deep-learning model. Users can customize the types of atom displayed (represented as circles with atom numbers). Curved lines connect atoms with predicted or measured NOE cross peaks; in this instance, the lines represent predicted cross peaks that have been validated through use of the slider tool. The atom circles are color-coded based on their assignment status: gray circles indicate unassigned atoms, while other colors (dark-green, light-green, orange, red) reflect the degree of agreement between the assigned and predicted chemical shift values. It is important to note that orange and red atom colorings do not necessarily indicate incorrect assignments, but they may warrant increased scrutiny. **(b)** Clicking on an atom circle opens a pop-up window displaying information about the chemical shift prediction for that atom, exemplified here for atom 6.H1*^′^*. Top row: measured versus predicted chemical shift values. Middle row: attributes input into the prediction model, which considers information about the base of interest, and the two closest bases on either side, along with H-bonded partners. For the bases two away, only information regarding whether the base features a purine (P) or pyrimidine (p) is provided. Bottom row: Information about examples present in the training dataset with the identical set of attributes as the atom of interest, including the number of examples, and the mean, standard deviation and full range of their chemical shifts.

Simulated peaklists were then generated based on the predicted correlations in each of the experiments. Several peak-box generation routines are available in NMRFx for various types of experiments. For the ^1^H-^1^H NOESY we used a built-in method that uses the predicted secondary structure alongside a distance database. The database was assembled by scanning a library of 3D RNA structures downloaded from the PDB, categorizing residues by secondary structure type, and measuring distances between atoms. Given the secondary structure of a novel RNA, predictions are made for atom pairs with occurrences in the database that would give rise to NOE cross-peaks.

The ^1^H-^1^H NOESY, ^1^H-^1^H TOCSY and ^1^H-^13^C HMQC spectra each contain several regions of interest. The layout tool was employed to display the aromatic-H1*^′^* and H1*^′^*-H1*^′^* regions of the NOESY and TOCSY spectra, along with the aromatic and H1*^′^*-C1*^′^* regions of the HMQC spectrum in a compact fashion, as shown in Figure 10. The peak slider tool was then activated to perform the assignments. Despite differences between the predicted chemical shifts and the spectrum, as seen in Figure 11, the predictions were robust enough to facilitate several initial assignments, particularly for outliers such as 33.H2, 11.H2, 7.H8 and 20.H8. These initial assignments served as a foundation, enabling the assignment of most aromatic and H1*^′^* chemical shifts in under a day with high confidence. Figure 10 illustrates the project’s status post-analysis, with red peak boxes indicating frozen assignments. Further experiments would be necessary to resolve uncertainties in the remaining ribose chemical shifts. It is straightforward to retrospectively add additional datasets to the project and expand the peak network to facilitate further assignment.

**Figure 10:**
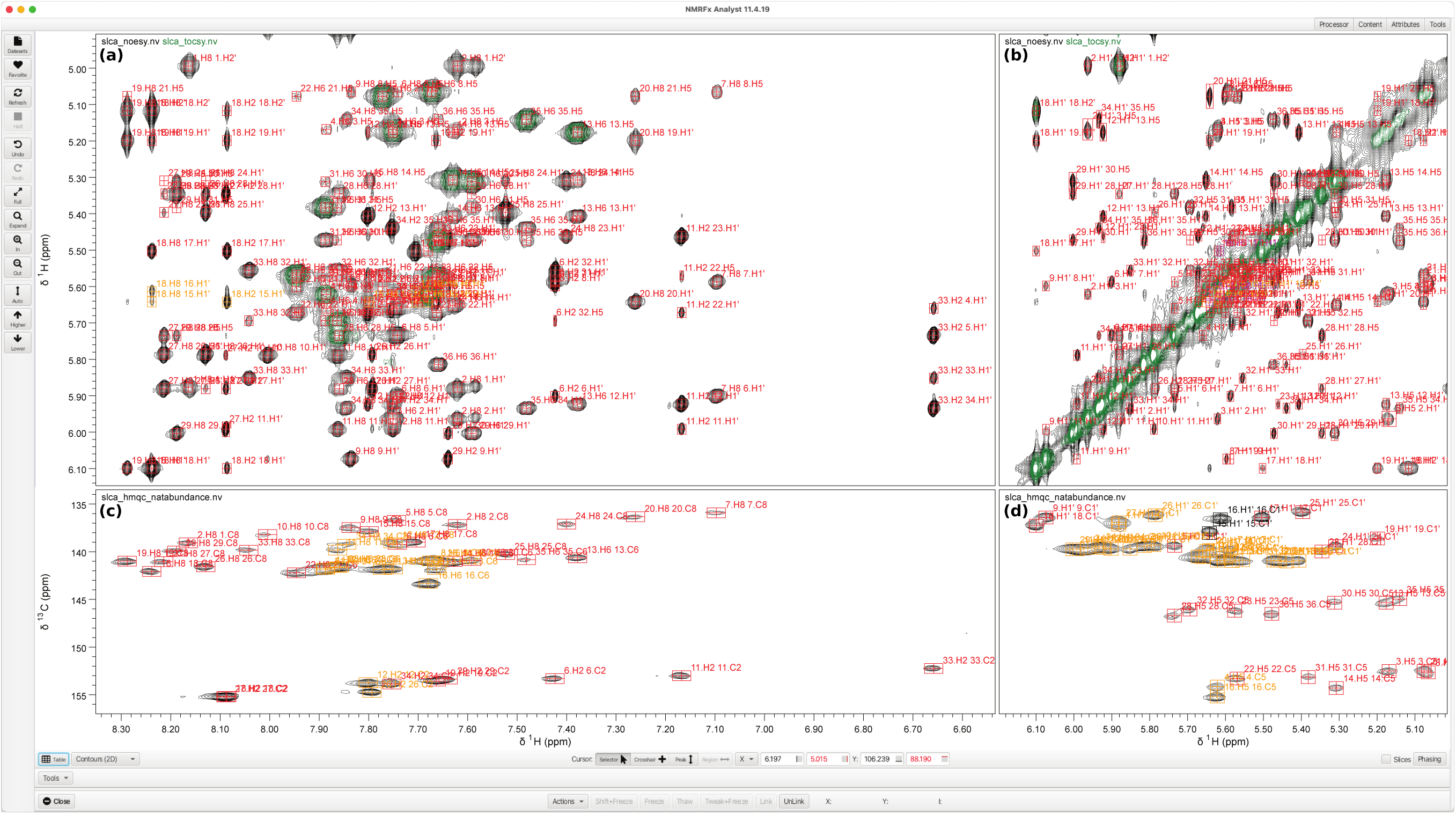
The use of NMRFx’s layout capability to visualize spectra related to the 36 nt RNA construct. **(a), (b)** Displays of both the ^1^H-^1^H NOESY (black) and ^1^H-^1^H TOCSY (green) spectra, with the visible region configured to show peaks associated with (a) aromatic-H1*^′^* and (b) H1*^′^*-H1*^′^* correlations. **(c), (d)** The ^1^H-^13^C HMQC spectrum, with the visible configured to display peaks corresponding to (c) aromatic and (d) H1*^′^*-C1*^′^* correlations. The charts are synchronized, such that when the user navigates one spectrum, the linked axes in the other spectra are automatically updated. The compact display of multiple regions of interest can be made advantage of when the peak slider tool is employed (see also Figure 11). The figure shows an advanced stage in the assignment process using the slider tool, with a majority of the assignments completed (red boxes).

**Figure 11:**
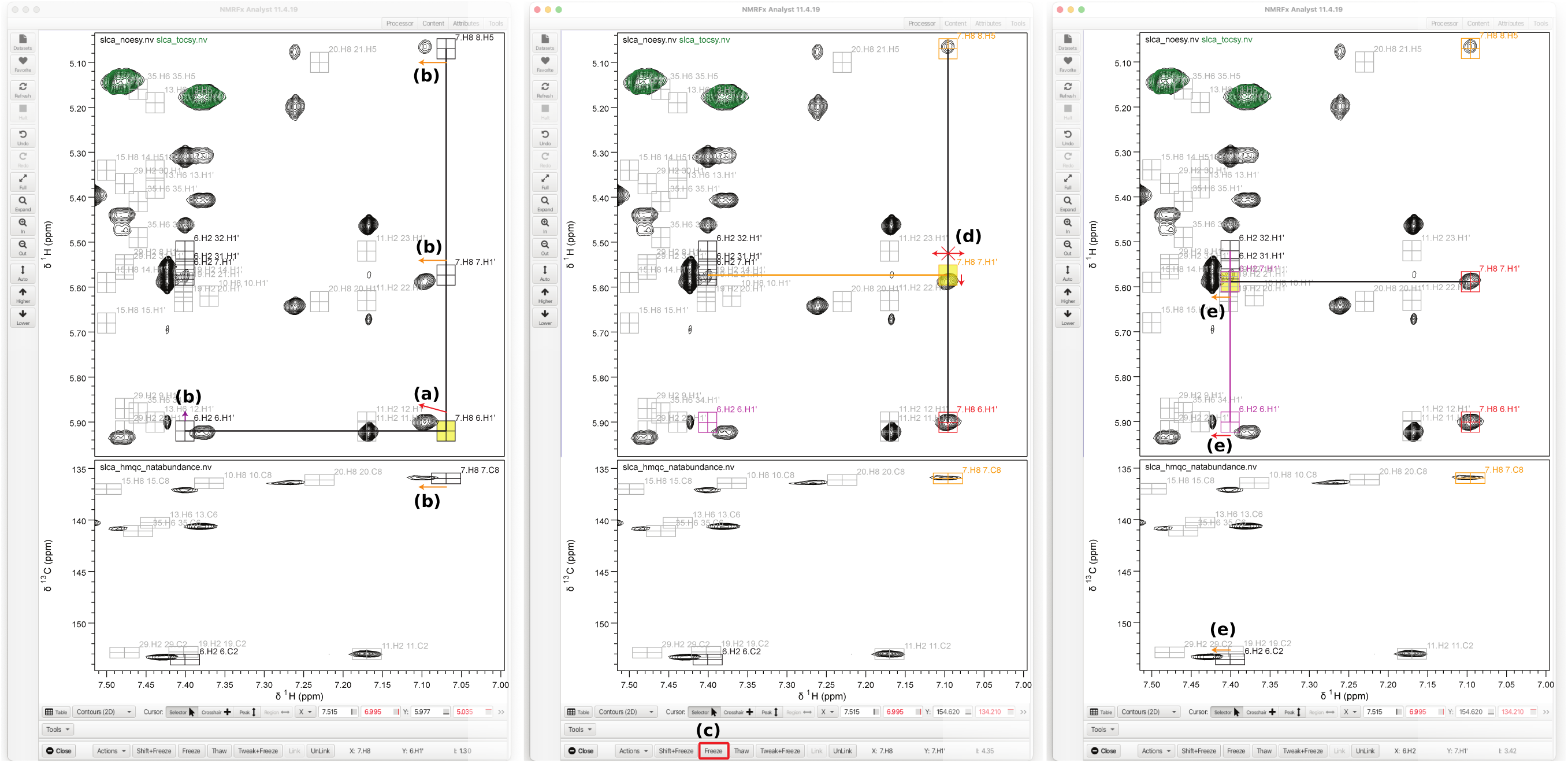
The peak slider tool being used to assign peaks associated with the 36 nt RNA construct. In the top chart, both the ^1^H-^1^H NOESY (black) and ^1^H-^1^H TOCSY (green) spectra are displayed, while the bottom chart shows the ^1^H-^13^C HMQC spectrum. **(a)** The peak-box corresponding to the predicted NOE between 7.H8 and 6.H1*^′^* is interactively adjusted based on close agreement with a peak in the observed NOESY spectrum. **(b)** Peak-boxes sharing an assignment are simultaneously updated. **(c)** Good agreement across the network of linked peaks suggests correct placement of this peak-box, which is therefore frozen in place by clicking the Freeze button. **(d)** Linked peak-boxes can no longer be adjusted in the frozen dimensions, but still can be in other dimensions, which is indicated by purple and orange colorings, indicating whether the peak can be moved in the *x*- or *y*-axis, respectively. **(e)** Assignment of 6.H2 is facilitated by two partially frozen peak-boxes which can only be adjusted in the *x*-axis.

After assigning the spectra, it is possible to generate distance and dihedral angle restraints and residual dipolar couplings (RDC) to facilitate a 3D structure calculation. NMRFx does not currently support the calculation of the gradients of RDC values. Instead, an optimizer that does not use gradients (CMA-ES [74]) is used to minimize the deviation of predicted RDCs (using a fit based on Singular Value Decomposition) from the experimental values. Although slower than using a gradient-based optimizer, it allows structures to be optimized against any criterion for which a numerical ranking can be made. Figure 12 shows one of the resulting structures and a corresponding plot of the predicted and experimental RDC values. Summary information on violations for the ten structures with lowest target function is available in the Supplementary Material.

**Figure 12:**
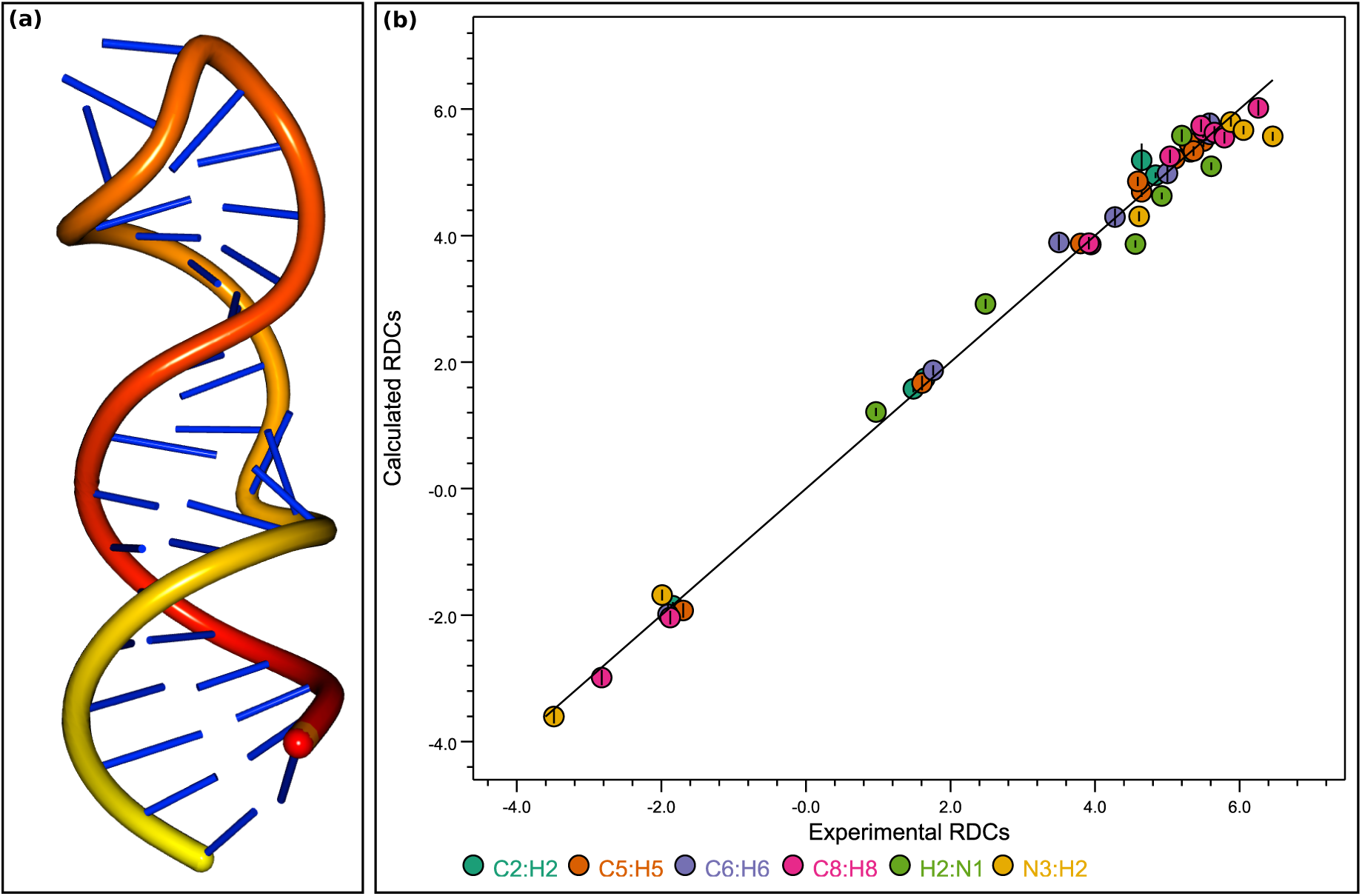
The result of performing a 3D structure calculation on the 36 nt RNA construct, based on distance, angle and RDC ([72] constraints determined by NMR assignment. **(a)** The structure within the computed 50-member ensemble with the lowest RMSD between the experimental and calculated RDC values. **(b)** A plot of normalized experimental versus estimated RDC values for the structure. The RMSD between the points and the black line, which defines equality between the experimental and calculated RDCs, is 0.04.

## 4 Conclusions

NMRFx is an integrated application that provides access to a wide range of features, making it a valuable means of analyzing NMR data in structural biology and beyond, as highlighted by its application in four diverse case-studies. The Java implementation allows cross-platform compatibility, and enables efficient use of multi-core hardware. Support for Python and Java extensions (including plugins) allows users to extend this open-source program with new capabilities, and its support for standardized file formats allow it to be used in conjunction with other applications. NMRFx is under active development and we expect that its existing and forthcoming tools, especially those supported by its integrated deep learning capabilities, will facilitate protocols using both standard and novel analysis methods.

## 5 Methods

### 5.1 NUS Processing Example

The spectrum presented in Figure 1 was generated by processing one of the datasets used in the NUScon competition [75]. The NUScon datasets were constructed by injecting synthetic peaks into experimentally acquired, uniformly-sampled NMR datasets, and subsequently applying exponentially-biased sampling schedules to form NUS datasets. The intensity and extent of overlap exhibited of the synthetic peaks, and the sampling scheme applied, were varied across the datasets to allow for different degrees of difficulty. Tables 2 and 4 of the NUScon publication [75] provide an outline of all the NUS datasets. The dataset processed and presented in Figure 1 is the HNCACB dataset of Protein A, with synthetic peak table 1, and 8.8% sampling. The data was provided as a series of nmrPipe time-domain files of the form test%03d.fid, along with a schedule file nuslist 3.scd, which NMRFx is able to import and process.

### 5.2 Chemical Shift Prediction

To train the protein shift prediction model, a dataset of 1480 non-paramagnetic proteins comprising standard residues with matching BMRB and PDB entries was used. For testing, a dataset of 61 proteins previously used for ShiftX2 was used [52]. The prediction results as calculated with the test proteins are listed in Table 3 of the Supplementary Information. Table 4 of the Supplementary Information lists the values of the attributes used for prediction of chemical shifts for glutamine residues as an example. Similar attribute sets with varying values are derived for each atom and amino acid type.

The RNA chemical shift prediction was trained with data assembled as previously described [58]. The current dataset consists of 371 RNA molecules that have assignments in the BMRB and associated PDB structures. Attributes comprising of secondary structure motifs (loops, bulges, commonly occurring tetraloops) and nucleotide base pairing were extracted and organized into training files. Training was performed with factorization machine (FM) regression model optimized with stochastic gradient descent as implemented in the Tribuo library (https://tribuo.org). Supplementary Table 5 describes model evaluation by cross-validated rms deviation for each atom group (^1^H, ^13^C, ^15^N) and provides a comparison with previously implemented support vector regression models [58]. The updated FM model robustly handles a significant increase in data points while maintaining high performance.

### 5.3 Structure Calculation of the 100 Protein Dataset

For each protein in the 100 protein structure dataset, a NEF file containing NMR constraints was processed by NMRFx using the following command-line program:

~~~
nmrfxs batch -n 200 -k 10 -a <nef-file>
~~~

where `<nef-file>` is a placeholder for the NEF file. The command generates an ensemble of 200 structures `(-n 200`), retains the ten with the lowest target function value (`-k 10`), and aligns the structures (`-a`). The output is a single PDBx/mmCIF file containing the structures of the refined 10-structure ensemble. No alterations to the method was made to optimize the calculation for any individual protein. A summary of violations is generated for each protein and an example of this is shown in Supplemntarty Table 6.

### 5.4 Algae Metabolomics Data Processing

The green alga *Chlorella vulgaris* UTEX 395 was used in this study and the details of the culturing conditions and the ^13^C labeling strategy as well as the data acquisition via High-Resolution Magic Angle Spinning can be found in the Supplementary Information. The resulting FIDs were imported into NMRFx using a tab-separated table containing the path to the data, the time of data collection, the replicate number, and whether the data were ^13^C coupled or decoupled. Using the Scanner Tool, the last decoupled dataset (the one with the strongest signals) was apodized with 0.3 Hz of line broadening, zero filled, Fourier transformed and phased. The baseline was corrected using manually selected regions and a third-order polynomial. All of the spectra in the table were then processed using those parameters and the spectra displayed in stacked mode. The spectra were subsequently aligned using the TMSP peak at 0 ppm. The spectra were not normalized because the cell pellets were of uniform cell count and the triacylglyceride accumulation mirrored quantification obtained by gas chromatography fatty acid methyl ester (GC-FAME) anaylysis. Integral regions were defined (i.e. for the methyl region between 0.8508 ppm to 0.9457 ppm) for each of the distinct regions of the fatty acid resonances Figure 8 (d) which creates a new column for each spectrum (row) in the Scanner Tool. The resulting table of integrals can then be exported as tab or comma separated data and imported into any external program for calculation of percent ^13^C incorporation at any given site as a function of time. The data can also be plotted within NMRFx using the built-in plotting and fitting tools.

### 5.5 Taccalonolide E Resonance Assignments

The resonance assignments for taccalonolide E were made using 1-dimensional ^1^H and ^13^C spectra as well as 2-dimensional HSQC, TOCSY, HMBC and ROESY datasets. Details of the sample preparation, data acquisition, and processing can be found in the Supplementary Information section. Reference chemical shift predictions were generated using PubChem file Conformer3D COMPOUND CID 56672430.sdf. The Peak Slider tool was used to match the predicted chemical shifts and predicted spin system patterns to the experimental data. The Analyze function within NMRFx was used to peak pick, integrate and define the multiplet structure for the 1-dimensional ^1^H spectrum and to produce the report in J. Org. Chem. format.

## Acknowledgments

B.A.J. would like to thank the many research groups who have provided data, made suggestions, and asked questions that helped us improve NMRFx. And special thanks are due to Jannalie Taylor and Vincent Fiack for help with software engineering.

S.G.H. would like to thank Prof. Adam Schuyler, UConn Health, for useful discussions regarding the NUScon datasets.

G.L.H. would like to thank Susan Mooberry and April Risinger, University of Texas Health San Antonio, for providing the sample of taccalonolide E and Rob Gardner (deceased), Bill Hiscox, Washington State University, Todd Pedersen, Brent Peyton and Robin Gerlach, Montana State University for assistance with growing the algae cultures and acquiring HRMAS data. G.L.H. would also like to thank the Washington State University Center for NMR Spectroscopy for providing instrument time to acquire data for both the taccalonolide E and algal metabolomics projects.

This work was supported in part by grants from the National Institute of General Medical Sciences of the National Institutes of Health, R01 GM123012 and RM1 GM145397 to B.A.J., and the National Institute of Allergy and Infectious Diseases of the National Institutes of Health, U54 AI170660 to B.A.J., J.M., and M.F.S., and R01 AI150498 to M.F.S., and from the Howard Hughes Medical Institute to M.F.S.

The content is solely the responsibility of the authors and does not necessarily represent the official views of the National Institutes of Health.

## Author Contributions

**Conceptualization:** B.A.J. **Methodology:** B.A.J. **Software:** E.K., S.G.H., K.M.C., B.A.J. **Validation:** E.K., S.G.H., G.L.H., M.F.S., J.M., B.A.J. **Formal Analysis:** E.K., S.G.H., B.A.J., J.M. **Investigation:** E.K., S.G.H., G.L.H., J.M., B.A.J. **Data Curation:** E.K., S.G.H., G.L.H., J.M., B.A.J. **Resources:** G.L.H., M.F.S., J.M., B.A.J. **Writing - Original Draft:** G.L.H.,E.K., S.G.H., J.M., B.A.J. **Writing - Review & Editing:** G.L.H., E.K., S.G.H., J.M., M.F.S., B.A.J. **Visualization:** E.K., S.G.H., B.A.J., J.M. **Supervision:** B.A.J. **Project Administration:** B.A.J. **Funding Acquisition:** M.F.S., J.M., B.A.J.

## Code availability

The NMRFx Analyst source code is available under the GNU General Public License, v3.0 at https://github.com/nanalysis/nmrfx. Executable versions of the software are available at http://nmrfx.org. Extensive documentation is provided at https://docs.nmrfx.org/.

