## Supplemental Information for "NMRFx: Integrated Software for NMR Data Processing, Visualization, Analysis and Structure Calculation"

August 22, 2025

**2 Sample, Data Acquisition and Processing** **5**

**3 Algae metabolomics** **6**

**4 Tables** **8**

**5 Code Listings** **19**

**References** **24**

### Acronyms

**BMRB** Biological Magnetic Resonance Data Bank

**CYANA** combined assignment and dynamics algorithm for NMR applications

**FM** factorization machine

**LARS** Least Angle Regression

**MAE** mean absolute error

**NMR** nuclear magnetic resonance spectroscopy

**PDB** Protein Data Bank

**RMSD** root-mean-squared deviation

**RNA** ribonucleic acid

**SVR** support vector regression

### 1 Futher Information on NMRFx Features

#### 1.1 NMRFx project structure

Project management is performed in NMRFx with two primary features. First, the state of a project is saved in a project directory. The project directory contains three primary components. The "datasets" subdirectory contains hard links to NMR datasets (typically resulting from processing in NMRFx. The linking method allows the user to have multiple projects that use the same datasets, but with only a single copy of the data rather than one copy in each project. The "windows" subdirectory contains one or more files (in YAML format) that describe the contents of the spectrum display windows. In this way, when the application is restarted and a project is reloaded, the exact display of the windows present when the project was saved is recreated. Finally, the "star" subdirectory contains a file in the NMR-STAR format used by the BMRB [1]. The NMR-STAR format is used by NMRFx to store the topology (sequence etc.) of the molecule under investigation and all derived data including peaks and chemical shifts.

#### 1.2 NMR Data Formats

NMRFx can read datasets in a variety of vendor formats. These include Bruker, Varian/Agilent, Jeol (partial functionality currently), SpinIT, the JCAMP format used by a variety of vendors, and some files used by the nmrPipe software. All formats are read directly, rather than being converted to an intermediate NMRFx format. Processed data can be written in the NMRFx Analyst format (common with that previously used by NMRViewJ), the UCSF format (used by Sparky and POKY), nmrPipe and SpinIT. Processed data files in formats used by NMRFx, UCSF Sparky, nmrPipe, Bruker, and JCAMP can be read for visualization and analysis.

##### 1.2.1 BMRB NMR-STAR

As mentioned above, the BMRB NMR-STAR format is used as the primary method for storing project data. Many data attributes of an NMR-STAR file are accessible in NMRFx (molecular topology, chemical shifts, peak lists etc.). Data that is not directly used is still loaded and stored internally and is accessible through custom scripts. We have worked with BMRB staff to understand and use their API to access the BMRB. Users can now search for data from the BMRB within NMRFx Analyst with the result displayed as a table of found entries. A selected entry can be directly fetched from the BMRB. If no project is currently active, all the contents of the entry are loaded. If a project is active, the user can select the data to load. For example, a user may have a molecular topology (protein or RNA sequence, for example), datasets, and peak lists, and retrieve only the chemical shifts from the BMRB entry.

Many journals require that chemical shift assignments for macromolecules be deposited in the BMRB prior to publication. NMRFx Analyst now facilitates this process by allowing the direct upload of an NMR-STAR file from an NMRFx Analyst project to the BMRB. The user interface dialog for uploads prompts the user to specify what categories of data should be included, whether to perform a test or production upload, and their email address. After upload, you will receive an email with a link to the BMRB site where you can complete any additional requirements. Traditionally, NMR users typically only upload a minimal set of required information to the BMRB (molecular topology and chemical shift assignments).

We hope that this direct upload tool will facilitate the deposition of more categories of data (peak lists, peak paths from titrations, relaxation data, etc.) and thereby make more project information accessible for validation, downstream analysis, and model training [2].

###### 1.2.2 NEF (NMR Exchange Format)

The NMR Exchange Format [3] has been designed as a method to provide NMR restraint data in a format that can be read by multiple structure calculation programs. Although NMR restraints can be represented in NMR-STAR files, a concern was that the high complexity of NMR-STAR (since it is designed for complete database depositions) was a barrier to developers of structure calculation programs. Depositions of structures in the wwPDB should be accompanied by NEF files that contain the molecular topology and restrictions.

NMRFx Analyst supports reading and writing NEF files. This can be used to initiate a project by reading a NEF file containing relevant information, for creating depositions to the wwPDB, and for structure calculations. For example, the structure module of NMRFx Analyst (vide infra) can calculate a three-dimensional model with a command as simple as "nmrfxs gen file.nef" where file.nef contains the topology and restraints.

###### 1.2.3 PDB and mmCIF

Basic reading and writing of PDB and mmCIF [4] files are supported. A molecular topology can be initiated from PDB files in two ways. In the default method, the PDB file is parsed to generate the amino-acid or nucleotide sequence of the macromolecule, and a residue library (based on rigid geometry concepts [5]), within NMRFx Analyst is referenced to construct atoms and bonds for each residue. This ensures that complete residues, with hydrogen atoms, are generated even when the PDB file is missing atoms. Atoms with entries in the file have their coordinates set based on the values in the file. The coordinates of missing atoms are generated, if possible, based on information from the residue library and the internal coordinates. This way, hydrogen atoms not present in the PDB file have appropriate coordinates generated. In the second method, the PDB file is read directly and the internal topology contains only those atoms in the file.

###### 1.2.4 Small Molecule Files

Small molecule structures that could act as ligands in a macromolecular project, or in their own right as in a medicinal chemistry project, can be read from several standard formats. These include `.mol`, `.sdf`, and `.mol2` files. In addition, SMILES [6] strings can be parsed to generate small molecule structures.

#### 2 Sample, Data Acquisition and Processing

A total of 7 mg of taccalonolide E was dissolved into 0.6 mL of $CDCl_3$ (Cambridge Isotope Labs, Woburn, MA) and placed into a Wilmad 535pp 5 mm NMR tube (SP-Wilmad Labglass, Vineland, NJ). Data were collected on either a Varian Inova 500 (Varian Associates, Palo Alto, CA) operating at 499.9 MHz for $^1$H and 125.7 MHz for $^{13}$C or a Varian VNMRS 600 operating at 599.7 MHz and 150 MHz for $^1$H and $^{13}$C respectively. 1-dimensional $^1$H spectra (600 MHz) were acquired with a 5020 Hz spectral width, a 45° pulsewidth, 5s relaxation delay and 2.5 s acquisition time. Sixteen transients were accumulated and the FID apodized with

0.3 Hz of exponential line broadening, zero filled and Fourier transformed. The 1-dimensional $^{1}$H spectrum was subsequently integrated, peak picked and the multiplets defined using the 'Analyze' function within NMRFx. 1-dimensional $^{13}$C spectra (150 MHz) were acquired with a 36,765 Hz spectral width, a 45 degree pulsewidth, 0.5 s relaxation delay and a 1.3 s acquisition time using gated broadband $^{1}$H decoupling. A total of 320 transients were accumulated and the FID apodized with 0.5 Hz of line broadening, zero filled and Fourier transformed. Two-dimensional HSQC spectra were acquired (600 MHz) using the Varian GHSQCAD pulse program and echo-antiecho for phase discrimination in F1. Spectral widths were 5020 Hz and 23,371 Hz and 0.23 s and 0.011 s acquisition times in the $^{1}$H and $^{13}$C dimensions respectively. A total of 256 increments using 16 scans per increment were acquired using uniform-sampling. The data were processed using 90 degree shifted sine bell squared apodization in both dimensions, zero filled and the indirect dimension was doubled in length using the NESTA algorithm prior to Fourier transform. Two-dimensional TOCSY spectra were acquired (600 MHz) using the Varian userlib Ztocsy_zq pulse program [7] and States-TPPI for phase discrimination in F1. Spectral widths were 5020 Hz in both dimensions and acquisition times were 0.204 s and 0.025 s in the direct and indirect dimensions respectively. A total of 256 increments using 4 scans per increment were acquired using uniform-sampling. The data were processed using 90 degree shifted sine bell squared apodization in both dimensions, zero filled and the indirect dimension doubled in length using the NESTA algorithm prior to Fourier transform. The spin lock for TOCSY transfer was 70 ms in duration using a DIPSI-2 mixing sequence. Two-dimensional ROESY spectra were acquired (500 MHz) using the Varian ROESY pulse program and States-TPPI for phase discrimination in F1. Spectral widths were 5006 Hz in both dimensions and the acquisition times were 0.204 s and 0.02 s in the direct and indirect dimensions respectively. Two hundred increments (uniform-sampling) were collected using 16 scans per increment and the data processed in a similar fashion to the TOCSY data. A 200 ms spinlock using the transverse Roesy mixing scheme [8] was used to effect cross relaxation. Two-dimensional HMBC spectra were acquired with the Varian `gHMBC` pulse program with spectral widths of 5102 and 39,912 Hz with acquisition times of 0.128 and 0.0065 s in the $^{1}$H and $^{13}$C dimensions respectively. A total of 400 increments using 64 scans per increment were acquired using uniform-sampling. A long range correlation delay (0.0625 s) corresponding to an 8 Hz long-range J value was used and a two-step, 1-bond J suppression sequence was used to lessen the 1-bond $^{13}$C-$^{1}$H correlations. The data were processed with unshifted sine bell apodization, zero filled and the indirect dimension doubled using the NESTA algorithm prior to Fourier transform.

### 3 Algae metabolomics

*Chlorella vulgaris* UTEX 395 was cultured using the methods described in [9] where $NaH^{13}CO_3$ (Cambridge Isotope Laboratories, Woburn, MA) was substituted for the unlabeled bicarbonate. The culture was grown for 38 h, and 1.5 mL aliquots were taken at various time points. The aliquots were centrifuged for 1 min at 5000 ×g, and the pellets were suspended in 34 µL of $D_2O$ containing 0.1 mg mL$^{-1}$ of TMSP as an internal standard. The suspended cells were transferred to a 4 mm glass rotor for analysis by high-resolution magic angle spinning (HRMAS) $^{1}$H NMR spectroscopy. The HRMAS spectra were acquired on a 600 MHz Varian VNMRS spectrometer ($^{1}$H and $^{13}$C carrier frequencies of 599.7 MHz and 150.8 MHz, respectively), spinning at a rate of 2.7 kHz, and oriented at the magic angle. The spectral width was 8000 Hz with an acquisition time of 0.4 s. The relaxation delay was 3.5 s during which time the residual $H_2O$ was saturated with a weak RF field of 50 Hz. $^{13}$C decoupling was accomplished using a GARP-1 modulated sequence with the decoupler offset placed at 60 ppm. $^{1}$H spectra were acquired in an interleaved fashion to minimize heating effects from the heteronuclear decoupling with 16 scans without $^{13}$C decoupling followed by 16 scans with $^{13}$C decoupling until 128 scans were acquired for both modes. The coupled and decoupled FIDs for each time point were separated into two files denoted by the string nnn for coupled and nny for decoupled.

### 4 Tables

Table 1: Evaluation of structure recalculations by NMRFx and CYANA [10] using deposited restraints in the 100-Protein NMR Spectra dataset [11]. RMSD values were calculated by superimposing the backbone (i.e. N, C, $C^{\alpha}$) atoms of the mean calculated structure and the mean PDB structure, using NMRFx's command-line program `super`. PDB IDs labeled with an asterisk do not have an associated CYANA recalculation.

| | | | RMSD (Å) | |
|---|---|---|---|---|
| | PDB ID | Residue ranges | CYANA | NMRFx |
| 1 | 6SVC* | 7–29 | – | – |
| 2 | 2JVD | 4–37 | 0.43 | 0.33 |
| 3 | 2K57 | 5–52 | 0.71 | 0.70 |
| 4 | 6SOW | 8–55 | 0.48 | 0.49 |
| 5 | 2LX7 | 5–59 | 1.02 | 1.27 |
| 6 | 2MA6 | 10–57 | 1.02 | 1.15 |
| 7 | 2JRM | 6–47 | 0.61 | 0.67 |
| 8 | 1YEZ | 15–25, 29–66 | 0.76 | 0.84 |
| 9 | 2L9R | 13–56 | 0.55 | 0.49 |
| 10 | 2K52 | 7–70 | 0.94 | 0.99 |
| 11 | 2KRS | 2–61 | 0.49 | 0.62 |
| 12 | 2K53 | 8–28, 39–66 | 0.69 | 0.49 |
| 13 | 2JT1 | 5–57, 66–69 | 0.72 | 0.78 |
| 14 | 2JVO | 6–71 | 0.76 | 0.71 |
| 15 | 2ERR | 2–75 | 1.02 | 1.11 |
| 16 | 2L1P | 19–78 | 1.29 | 1.69 |
| 17 | 2LN3 | 6–72 | 0.56 | 0.77 |
| 18 | 2HEQ | 17–20, 34–68 | 0.65 | 0.71 |
| 19 | 2KK8 | 10–82 | 1.12 | 1.16 |
| 20 | 2KD0 | 13–81 | 1.01 | 1.21 |
| 21 | 2LML | 3–78 | 1.30 | 1.15 |
| 22 | 2K3D | 2–81 | 0.84 | 0.72 |
| 23 | 2LK2 | 14–65 | 0.95 | 1.01 |
| 24 | MH04* | – | – | – |
| 25 | 1PQX | 13–19, 28–34, 38–65, 70–81 | 1.14 | 0.98 |
| 26 | 2L33 | 19–36, 46–79 | 0.43 | 0.44 |
| 27 | 2KZV | 9–80 | 1.17 | 1.24 |
| 28 | 2KCT | 11–37, 44–83 | 0.98 | 1.10 |
| 29 | 2MDR | 9–89 | 1.37 | 1.47 |
| 30 | 2FB7 | 20–52, 74–87 | 0.45 | 0.51 |
| 31 | 2MB0 | 8–41, 49–85 | 0.65 | 0.66 |
| 32 | 2L05 | 19–89 | 0.81 | 0.98 |
| 33 | 2KJR | 15–23, 28–94 | 0.69 | 0.77 |
| 34 | 2M5O | 17–91 | 0.75 | 1.11 |
| 35 | MDM2* | – | – | – |
| 36 | 2LNA | 16–49, 59–94 | 0.90 | 0.89 |

| | PDB ID | Residue ranges | RMSD (Å) | |
|---|---|---|---|---|
| | | | CYANA | NMRFx |
| 37 | 2LA6 | 15–97 | 0.79 | 0.69 |
| 38 | 6FIP | 11–93 | 0.86 | 0.91 |
| 39 | 2LEA | 15–45, 55–88 | 0.35 | 0.56 |
| 40 | 2LL8 | 4–90 | 0.94 | 1.04 |
| 41 | 2KPN | 12–84 | 0.83 | 0.67 |
| 42 | 2K0M | 7–70, 76–93 | 0.75 | 0.76 |
| 43 | 2K5V | 2–29, 38–78, 83–94 | 1.03 | 1.04 |
| 44 | 2MQL | 15–84 | 0.53 | 0.64 |
| 45 | 2K75 | 3–92 | 0.91 | 1.14 |
| 46 | 2LTM | 14–99 | 0.81 | 0.73 |
| 47 | 2KOB | 3–93 | 0.96 | 0.63 |
| 48 | 2KHD | 31–97 | 1.15 | 1.40 |
| 49 | 2RN7 | 10–55 | 0.51 | 0.59 |
| 50 | 2LXU | 9–95 | 0.73 | 1.41 |
| 51 | 2KIF | 3–97 | 0.72 | 0.73 |
| 52 | 2KBN | 5–29, 34–54, 58–76, 81–94 | 0.40 | 0.62 |
| 53 | 2MK2 | 14–108 | 1.17 | 1.42 |
| 54 | 2K50 | 10–34, 42–85, 92–105 | 0.93 | 0.83 |
| 55 | 2KL5 | 12–53, 58–66, 76–86, 93–99 | 0.68 | 0.64 |
| 56 | 2LTA | 4–98 | 1.00 | 1.21 |
| 57 | 2KIW | 2–81 | 1.63 | 1.29 |
| 58 | 2LVB | 3–102 | 0.79 | 0.84 |
| 59 | 2LND | 3–48, 52–101 | 0.69 | 0.76 |
| 60 | 1WQU | 8–106 | 0.49 | 0.61 |
| 61 | 2KL6 | 6–106 | 0.85 | 0.73 |
| 62 | 6GT7 | 7–30, 40–87, 94–113 | 0.56 | 0.63 |
| 63 | 2JN8 | 12–26, 31–109 | 0.78 | 0.87 |
| 64 | 2K5D | 19–50, 55–84, 98–107 | 0.70 | 0.76 |
| 65 | 2KD1 | 7–89 | 1.23 | 0.99 |
| 66 | 2LTL | 19–35, 39–41, 46–110 | 1.12 | 0.95 |
| 67 | 2KVO | 3–23, 28–103 | 0.72 | 0.89 |
| 68 | 1T0Y | 4–83 | 0.88 | 0.74 |
| 69 | 2KCD | 3–108 | 0.99 | 1.65 |
| 70 | 2KRT | 6–114 | 1.30 | 1.24 |
| 71 | 2LFI | 2–104 | 1.03 | 2.16 |
| 72 | 2JQN | 3–111 | 1.18 | 1.98 |
| 73 | 2L7Q | 12–37, 46–101, 105–114 | 0.76 | 1.31 |
| 74 | 2KFP | 3–115 | 0.71 | 0.81 |
| 75 | 1SE9* | 17–84, 94–101 | – | – |
| 76 | 2L3G | 13–123 | 0.65 | 0.74 |
| 77 | 2L3B | 14–38, 45–113 | 0.66 | 1.18 |
| 78 | 2LRH | 3–122 | 0.91 | 1.15 |
| 79 | 1VEE | 6–123 | 0.38 | 0.51 |
| 80 | 2K1G | 5–78, 83–122 | 0.77 | 0.69 |

| | | | RMSD (Å) | |
|---|---|---|---|---|
| | PDB ID | Residue ranges | CYANA | NMRFx |
| 81 | 2KKZ | 5–80, 86–118 | 1.17 | 1.14 |
| 82 | 1VDY | 9–102, 113–128 | 0.45 | 0.52 |
| 83 | 2KKL | 33–90, 96–125 | 0.50 | 0.89 |
| 84 | 2N4B* | 2–26, 40–54, 66–134 | – | – |
| 85 | 2L8V | 4–22, 37–65, 73–129 | 1.02 | 1.39 |
| 86 | 2LGH | 2–109, 113–135 | 1.37 | 1.22 |
| 87 | 2K1S | 3–140 | 0.94 | 1.74 |
| 88 | 2M4F | 23–46, 51–57, 63–94, 103–114, 120–129, 136–148 | 0.25 | 0.36 |
| 89 | 2JXP | 16–144 | 1.61 | 1.47 |
| 90 | 2L06 | 15–38, 45–141 | 0.77 | 1.10 |
| 91 | 2LAH | 14–25, 33–158 | 0.83 | 1.10 |
| 92 | 2LAK | 10–37, 68–77, 93–139 | 0.55 | 0.87 |
| 93 | 2L82 | 3–151 | 0.81 | 0.74 |
| 94 | 2M47 | 5–25, 40–56, 66–156 | 0.93 | 2.09 |
| 95 | 2K3A | 57–102, 108–127, 138–153 | 0.75 | 0.86 |
| 96 | 2M7U* | 12–151 | – | – |
| 97 | 2B3W | 16–162 | 1.01 | 1.07 |
| 98 | KRAS* | – | – | – |
| 99 | 2G0Q | 18–54, 60–126 | 0.91 | 1.16 |
| 100 | 2LF2 | 6–44, 53–69, 76–105, 111–165 | 1.24 | 1.67 |

Table 2: Smallest (Min) and largest (Max) distance violations across the 10 lowest energy structures generated by NMRFx structure recalculation for each protein in the 100-protein dataset (see Table 1). Values were not obtained for proteins whose PDB IDs are labeled with an asterisk.

| | | Distance (Å) | | | | Distance (Å) | |
|---|---|---|---|---|---|---|---|
| | PDB ID | Min | Max | | PDB ID | Min | Max |
| 1 | 6SVC* | – | – | 51 | 2KIF | 0.10 | 0.40 |
| 2 | 2JVD | 0.00 | 0.00 | 52 | 2KBN | 0.00 | 0.21 |
| 3 | 2K57 | 0.00 | 0.23 | 53 | 2MK2 | 0.00 | 0.22 |
| 4 | 6SOW | 0.31 | 0.45 | 54 | 2K50 | 0.17 | 0.42 |
| 5 | 2LX7 | 0.00 | 0.11 | 55 | 2KL5 | 0.11 | 0.53 |
| 6 | 2MA6 | 0.57 | 0.57 | 56 | 2LTA | 0.00 | 0.29 |
| 7 | 2JRM | 0.30 | 0.49 | 57 | 2KIW | 0.00 | 0.16 |
| 8 | 1YEZ | 0.00 | 0.29 | 58 | 2LVB | 0.12 | 0.27 |
| 9 | 2L9R | 0.00 | 0.14 | 59 | 2LND | 0.30 | 0.41 |
| 10 | 2K52 | 0.00 | 0.43 | 60 | 1WQU | 0.47 | 0.80 |
| 11 | 2KRS | 0.00 | 0.15 | 61 | 2KL6 | 0.21 | 0.28 |
| 12 | 2K53 | 0.30 | 0.83 | 62 | 6GT7 | 0.76 | 0.86 |
| 13 | 2JT1 | 0.00 | 0.31 | 63 | 2JN8 | 0.11 | 0.30 |
| 14 | 2JVO | 0.59 | 1.47 | 64 | 2K5D | 0.18 | 0.43 |
| 15 | 2ERR | 0.75 | 1.02 | 65 | 2KD1 | 0.22 | 0.59 |
| 16 | 2L1P | 0.00 | 0.22 | 66 | 2LTL | 0.00 | 0.24 |
| 17 | 2LN3 | 0.00 | 0.21 | 67 | 2KVO | 0.00 | 0.25 |
| 18 | 2HEQ | 0.00 | 0.41 | 68 | 1T0Y | 0.17 | 0.58 |
| 19 | 2KK8 | 0.00 | 0.20 | 69 | 2KCD | 0.00 | 0.29 |
| 20 | 2KD0 | 0.00 | 0.18 | 70 | 2KRT | 0.00 | 0.26 |
| 21 | 2LML | 0.00 | 0.33 | 71 | 2LFI | 0.14 | 0.49 |
| 22 | 2K3D | 0.00 | 0.36 | 72 | 2JQN | 0.00 | 0.36 |
| 23 | 2LK2 | 0.11 | 0.24 | 73 | 2L7Q | 0.00 | 0.41 |
| 24 | MH04* | – | – | 74 | 2KFP | 0.20 | 0.41 |
| 25 | 1PQX | 0.13 | 0.56 | 75 | 1SE9* | – | – |
| 26 | 2L33 | 0.11 | 0.39 | 76 | 2L3G | 0.00 | 0.27 |
| 27 | 2KZV | 0.00 | 0.23 | 77 | 2L3B | 0.00 | 0.25 |
| 28 | 2KCT | 0.00 | 0.28 | 78 | 2LRH | 0.14 | 0.48 |
| 29 | 2MDR | 1.56 | 1.60 | 79 | 1VEE | 0.61 | 1.01 |
| 30 | 2FB7 | 0.42 | 0.60 | 80 | 2K1G | 0.13 | 0.23 |
| 31 | 2MB0 | 0.39 | 0.70 | 81 | 2KKZ | 0.00 | 0.21 |
| 32 | 2L05 | 0.00 | 0.34 | 82 | 1VDY | 0.52 | 0.74 |
| 33 | 2KJR | 0.00 | 0.10 | 83 | 2KKL | 0.14 | 0.56 |
| 34 | 2M5O | 0.13 | 0.43 | 84 | 2N4B* | – | – |
| 35 | MDM2* | – | – | 85 | 2L8V | 0.00 | 0.16 |
| 36 | 2LNA | 0.18 | 0.37 | 86 | 2LGH | 0.73 | 0.76 |
| 37 | 2LA6 | 0.13 | 0.38 | 87 | 2K1S | 0.14 | 0.38 |
| 38 | 6FIP | 1.10 | 1.42 | 88 | 2M4F | 0.26 | 0.58 |
| 39 | 2LEA | 1.04 | 1.43 | 89 | 2JXP | 0.20 | 0.38 |

| | | Distance (Å) | | | | Distance (Å) | |
|---|---|---|---|---|---|---|---|
| | PDB ID | Min | Max | | PDB ID | Min | Max |
| 40 | 2LL8 | 0.00 | 0.37 | 90 | 2L06 | 0.00 | 0.39 |
| 41 | 2KPN | 0.13 | 0.28 | 91 | 2LAH | 0.34 | 0.76 |
| 42 | 2K0M | 0.11 | 0.18 | 92 | 2LAK | 0.14 | 0.23 |
| 43 | 2K5V | 0.00 | 0.21 | 93 | 2L82 | 0.12 | 0.49 |
| 44 | 2MQL | 0.14 | 0.41 | 94 | 2M47 | 0.15 | 0.42 |
| 45 | 2K75 | 0.00 | 0.19 | 95 | 2K3A | 0.37 | 0.49 |
| 46 | 2LTM | 0.00 | 0.51 | 96 | 2M7U* | – | – |
| 47 | 2KOB | 0.00 | 0.21 | 97 | 2B3W | 0.18 | 0.25 |
| 48 | 2KHD | 0.00 | 0.13 | 98 | KRAS* | – | – |
| 49 | 2RN7 | 0.00 | 0.13 | 99 | 2G0Q | 0.52 | 1.18 |
| 50 | 2LXU | 0.00 | 0.26 | 100 | 2LF2 | 0.00 | 0.16 |

Table 3: Evaluation of the LARS-based protein chemical shift predictions with ten-fold cross validation (CV) and on a test dataset comprising of 61 proteins, introduced in the paper outlining SHIFTX+ [12].

| Atom | CV RMSD (ppm) | Test RMSD (ppm) | Test MAE (ppm) |
|---|---|---|---|
| $^{1}H$ | 0.472 | 0.470 | 0.360 |
| $^{1}H^{\alpha}$ | 0.305 | 0.300 | 0.230 |
| $^{1}H^{\beta}$ | 0.256 | 0.250 | 0.190 |
| $^{1}H^{\gamma}$ | 0.231 | 0.230 | 0.170 |
| $^{1}H^{\delta}$ | 0.259 | 0.290 | 0.225 |
| $^{1}H^{\epsilon}$ | 0.315 | 0.327 | 0.250 |
| $^{1}H$ Methyl | 0.188 | 0.180 | 0.130 |
| $^{1}H$ Aromatic | 0.288 | 0.290 | 0.220 |
| $^{13}C$ | 1.304 | 1.070 | 0.840 |
| $^{13}C^{\alpha}$ | 1.16 | 0.870 | 0.670 |
| $^{13}C^{\beta}$ | 1.262 | 1.020 | 0.780 |
| $^{13}C^{\gamma}$ | 1.067 | 0.900 | 0.645 |
| $^{13}C^{\delta}$ | 2.028 | 0.620 | 0.460 |
| $^{13}C^{\epsilon}$ | 0.676 | 0.390 | 0.280 |
| $^{13}C$ Methyl | 1.315 | 1.170 | 0.910 |
| $^{13}C$ Aromatic | 1.236 | 1.120 | 0.820 |
| $^{15}N$ | 2.647 | 2.423 | 1.767 |

Table 4: Protein structure attributes and their coefficient values used by the LARS model to predict chemical shifts of the backbone atoms of glutamine. Other amino acids use the same attributes, but with different coefficient values. Attributes are sorted in descending order by the sum, over the atom types, of the absolute value of the coefficients, so more important attributes are near the top.

| Attribute | $^{13}C^{\alpha}$ | $^{13}C^{\beta}$ | $^{13}C$ | $^{15}N$ | $^{1}H$ | $^{1}H^{\alpha}$ |
|---|---|---|---|---|---|---|
| hydrogen shift1 | 0.0000 | 0.0000 | 0.0000 | 0.0652 | -1.6809 | 0.5948 |
| hydrogen shift3 | 0.0000 | 0.0000 | 0.0000 | -0.0176 | 1.7780 | -0.5381 |
| $\cos(\psi_i)$ | 0.1952 | -0.2550 | 0.0425 | -0.0886 | 0.3558 | -0.1756 |
| $\sin(\psi_i)\sin(\phi_i)$ | 0.1519 | -0.1201 | 0.0952 | -0.2172 | -0.0837 | -0.2804 |
| $\cos(\phi_i)$ | 0.1103 | -0.1775 | 0.0377 | 0.1650 | 0.1308 | -0.1833 |
| $\cos(\psi_{i-1})$ | -0.0039 | -0.0325 | -0.0244 | -0.2673 | -0.4064 | -0.0552 |
| $\sin(2\phi_i)$ | -0.1704 | 0.3059 | -0.0625 | -0.0539 | 0.0432 | 0.1113 |
| ring current shift | 0.0336 | 0.0881 | 0.0440 | 0.0021 | 0.2123 | 0.2356 |
| $\cos(3\psi_{i-1})$ | -0.0200 | 0.0189 | -0.0208 | 0.3886 | 0.0657 | 0.0754 |
| $\sin(\psi_i)$ | -0.1177 | -0.0967 | 0.0127 | -0.0578 | -0.1189 | -0.1786 |
| $\cos(\psi_i)\sin(\phi_i)$ | 0.0654 | -0.1504 | -0.0298 | -0.0521 | 0.0756 | -0.1164 |
| $\sin(2\psi_i)$ | -0.1099 | -0.1867 | -0.0841 | -0.0322 | -0.0405 | -0.0313 |
| $\cos(\phi_{i+1})$ | 0.0273 | -0.0916 | 0.1112 | 0.0027 | 0.0061 | -0.2196 |
| $\cos(\chi_i)$ | -0.0528 | -0.0877 | 0.0534 | -0.1073 | 0.1096 | 0.0357 |
| $\sin(\phi_i)$ | 0.0711 | -0.0706 | -0.0094 | -0.1707 | -0.0234 | -0.0988 |
| $\sin(\psi_{i-1})$ | 0.0061 | 0.0066 | -0.0581 | 0.1980 | 0.1215 | -0.0306 |
| $\cos(\psi_i)\cos(\phi_i)$ | -0.0700 | 0.0698 | -0.0201 | 0.0374 | 0.0470 | 0.1616 |
| $\sin(\chi_i)$ | -0.0646 | -0.0692 | -0.1045 | 0.0402 | 0.0800 | 0.0250 |
| $\cos(\psi_{i+1})$ | 0.0919 | -0.0704 | 0.1332 | 0.0529 | 0.0204 | -0.0013 |
| $\sin(\phi_i)\cdot \text{size}_{i-1}$ | -0.0305 | -0.0006 | -0.0300 | 0.2757 | 0.0267 | -0.0046 |
| $\cos(\chi_{2i})$ | -0.0972 | -0.0501 | -0.0189 | 0.0545 | -0.0837 | 0.0511 |
| $\cos(2\chi_{2i})$ | -0.0624 | 0.0739 | -0.0067 | -0.0295 | -0.1613 | 0.0107 |
| $\sin(\psi_i)\cdot \text{size}_{i+1}$ | -0.0321 | 0.0517 | -0.0306 | -0.0279 | 0.0701 | 0.1212 |
| $\cos(2\psi_{i-1})$ | 0.0139 | -0.0338 | -0.0196 | -0.2107 | -0.0436 | -0.0109 |
| $\cos(\phi_{i-1})$ | 0.0168 | -0.0854 | 0.0400 | -0.0738 | 0.0211 | -0.0863 |
| $\cos(\psi_i)\cdot \text{size}_{i+1}$ | 0.0070 | 0.1169 | 0.0005 | -0.0251 | -0.0747 | 0.0821 |
| $\sin(\chi_i)\sin(\chi_{2i})$ | -0.0259 | 0.0513 | 0.0304 | -0.0117 | -0.1731 | -0.0110 |
| $\cos(3\phi_{i-1})$ | 0.0292 | -0.0594 | -0.0171 | -0.0834 | 0.0841 | -0.0089 |
| $\cos(2\psi_i)$ | -0.0434 | 0.0485 | -0.0036 | -0.1289 | 0.0112 | -0.0420 |
| $\sin(\phi_i)\cdot \text{hydrophobicity}_{i-1}$ | -0.0348 | -0.0225 | -0.0255 | 0.1638 | 0.0023 | 0.0287 |
| $\sin(\phi_i)\cos(\chi_i)$ | -0.0906 | -0.0567 | 0.0194 | -0.0106 | 0.0435 | 0.0547 |
| $\cos(\chi_i)\cos(\chi_{2i})$ | 0.0505 | -0.0246 | -0.0064 | 0.0159 | 0.1313 | -0.0416 |
| $\cos(3\psi_i)$ | -0.0426 | 0.0273 | -0.0515 | -0.0804 | -0.0116 | 0.0401 |
| $\sin(\psi_{i+1})$ | -0.0584 | -0.0273 | -0.0320 | -0.0588 | -0.0475 | 0.0289 |
| $\cos(\phi_i)\cdot \text{size}_{i-1}$ | 0.0552 | -0.0664 | 0.0475 | -0.0484 | -0.0201 | -0.0052 |
| $\sin(\phi_i)\cdot \text{aromaticity}_{i-1}$ | -0.0112 | 0.0108 | -0.0211 | -0.0868 | -0.0860 | -0.0219 |
| $\sin(\psi_i)\sin(\chi_i)$ | 0.0311 | 0.0412 | 0.0373 | 0.0318 | 0.0566 | -0.0389 |
| $\sin(3\psi_i)$ | -0.0081 | 0.0755 | 0.0218 | -0.0636 | -0.0009 | 0.0633 |
| $\cos(2\phi_i)$ | -0.0232 | 0.0564 | -0.0247 | 0.0184 | 0.0639 | -0.0403 |
| $\sin(\psi_i)\sin(\psi_{i-1})$ | -0.0563 | 0.0191 | 0.0160 | -0.0333 | 0.0405 | 0.0544 |

| Attribute | $^{13}C^{\alpha}$ | $^{13}C^{\beta}$ | $^{13}C$ | $^{15}N$ | $^{1}H$ | $^{1}H^{\alpha}$ |
|---|---|---|---|---|---|---|
| $\sin(2\phi_{i+1})$ | 0.0569 | 0.0144 | 0.0430 | 0.0044 | 0.0845 | 0.0144 |
| $\cos(3\phi_{i})$ | -0.0240 | -0.0854 | 0.0153 | -0.0037 | 0.0256 | 0.0620 |
| $\cos(3\phi_{i+1})$ | 0.0024 | 0.0316 | -0.0499 | -0.0051 | -0.0853 | -0.0354 |
| $\cos(\psi_{i})\cos(\psi_{i-1})$ | -0.0156 | 0.0358 | 0.0206 | -0.0375 | 0.0570 | 0.0312 |
| $\cos(\psi_{i})\sin(\chi_{i})$ | 0.0306 | 0.0199 | 0.0258 | 0.0130 | 0.0872 | -0.0181 |
| $\sin(3\psi_{i-1})$ | 0.0150 | 0.0095 | 0.0204 | -0.0247 | -0.0629 | -0.0542 |
| $\sin(\psi_{i})\cdot \text{proline}_{i+1}$ | 0.0022 | 0.0827 | 0.0169 | -0.0475 | 0.0189 | -0.0101 |
| $\sin(\phi_{i})\cos(\chi_{2i})$ | -0.0292 | 0.0244 | -0.0023 | 0.0270 | -0.0453 | 0.0494 |
| $\sin(\phi_{i+1})$ | 0.0271 | 0.0326 | -0.0391 | 0.0142 | -0.0056 | -0.0589 |
| $\cos(\phi_{i})\cdot \text{hydrophobicity}_{i-1}$ | 0.0376 | -0.0516 | 0.0183 | -0.0358 | -0.0131 | -0.0197 |
| $\sin(\chi_{2i})$ | -0.0181 | 0.0105 | 0.0426 | -0.0113 | -0.0699 | 0.0225 |
| $\cos(3\psi_{i+1})$ | -0.0149 | 0.0384 | -0.0465 | 0.0286 | 0.0430 | 0.0024 |
| $\sin(\phi_{i})\sin(\chi_{i})$ | -0.0213 | -0.0230 | -0.0577 | 0.0014 | -0.0162 | -0.0509 |
| $\sin(\psi_{i})\cdot \text{hydrophobicity}_{i+1}$ | -0.0077 | 0.0063 | -0.0072 | -0.0494 | 0.0472 | 0.0469 |
| $\cos(\psi_{i})\cdot \text{proline}_{i+1}$ | -0.0244 | 0.0900 | 0.0071 | -0.0088 | 0.0023 | 0.0308 |
| $\sin(\phi_{i-1})$ | -0.0014 | 0.0262 | -0.0347 | 0.0077 | 0.0751 | -0.0125 |
| $\cos(2\chi_{i})$ | 0.0102 | 0.0413 | 0.0028 | 0.0101 | -0.0805 | -0.0097 |
| $\sin(\psi_{i})\cos(\phi_{i})$ | -0.0607 | -0.0297 | 0.0243 | -0.0261 | 0.0101 | -0.0019 |
| $\cos(\psi_{i})\cdot \text{hydrophobicity}_{i+1}$ | -0.0126 | 0.0349 | 0.0007 | -0.0187 | -0.0363 | 0.0480 |
| $\cos(2\phi_{i-1})$ | 0.0091 | -0.0536 | -0.0127 | -0.0115 | 0.0528 | 0.0105 |
| $\sin(\chi_{i})\cos(\chi_{2i})$ | -0.0121 | -0.0100 | 0.0059 | 0.0726 | 0.0398 | 0.0059 |
| $\sin(\phi_{i})\sin(\chi_{2i})$ | -0.0391 | -0.0045 | 0.0440 | 0.0116 | -0.0001 | 0.0417 |
| $\sin(\psi_{i})\cos(\psi_{i-1})$ | 0.0048 | 0.0021 | -0.0097 | -0.0527 | 0.0307 | -0.0389 |
| $\sin(2\psi_{i-1})$ | 0.0203 | -0.0220 | 0.0195 | 0.0124 | -0.0525 | -0.0110 |
| $\cos(\phi_{i})\cos(\chi_{2i})$ | -0.0062 | 0.0274 | 0.0027 | -0.0006 | 0.0563 | 0.0442 |
| $\sin(\phi_{i})\cdot \text{proline}_{i-1}$ | -0.0161 | 0.0192 | -0.0023 | 0.0367 | -0.0279 | 0.0317 |
| $\cos(2\psi_{i+1})$ | 0.0111 | -0.0422 | 0.0392 | 0.0081 | 0.0291 | -0.0034 |
| $\sin(3\phi_{i+1})$ | -0.0064 | 0.0236 | -0.0265 | -0.0260 | 0.0098 | -0.0351 |
| $\sin(3\phi_{i-1})$ | 0.0139 | -0.0039 | -0.0374 | 0.0294 | 0.0250 | -0.0114 |
| $\sin(2\chi_{2i})$ | 0.0407 | 0.0023 | -0.0005 | 0.0085 | 0.0422 | -0.0229 |
| $\cos(2\phi_{i+1})$ | -0.0291 | 0.0057 | 0.0048 | 0.0149 | -0.0124 | 0.0431 |
| $\sin(2\chi_{i})$ | 0.0182 | 0.0090 | 0.0085 | -0.0082 | -0.0370 | -0.0288 |
| $\cos(\phi_{i})\sin(\chi_{i})$ | -0.0110 | 0.0140 | 0.0011 | 0.0112 | -0.0260 | -0.0448 |
| $\sin(2\psi_{i+1})$ | -0.0441 | -0.0094 | -0.0424 | 0.0064 | -0.0020 | -0.0030 |
| $\sin(\psi_{i})\cos(\chi_{i})$ | 0.0215 | -0.0097 | -0.0289 | 0.0082 | 0.0165 | -0.0168 |
| $\sin(3\phi_{i})$ | -0.0578 | -0.0050 | 0.0251 | 0.0051 | -0.0052 | -0.0009 |
| $\cos(\psi_{i})\cos(\chi_{2i})$ | 0.0102 | 0.0389 | 0.0145 | -0.0018 | 0.0159 | 0.0156 |
| $\cos(\phi_{i})\cos(\chi_{i})$ | 0.0160 | 0.0134 | -0.0025 | 0.0237 | -0.0130 | -0.0281 |
| $\cos(\psi_{i})\cos(\chi_{i})$ | 0.0045 | -0.0078 | -0.0215 | 0.0259 | 0.0206 | 0.0114 |
| $\sin(\psi_{i})\cos(\chi_{2i})$ | 0.0028 | 0.0398 | -0.0009 | 0.0145 | -0.0027 | -0.0306 |
| $\cos(\psi_{i})\cdot \text{aromaticity}_{i+1}$ | 0.0020 | -0.0209 | -0.0044 | 0.0078 | 0.0517 | -0.0009 |
| $\sin(3\psi_{i+1})$ | 0.0043 | 0.0017 | -0.0279 | 0.0389 | -0.0004 | 0.0110 |
| $\cos(\phi_{i})\cdot \text{aromaticity}_{i-1}$ | -0.0058 | 0.0103 | -0.0150 | 0.0249 | 0.0020 | -0.0248 |
| $\cos(\psi_{i})\cdot \text{charge}_{i+1}$ | 0.0045 | 0.0127 | -0.0019 | 0.0183 | -0.0196 | 0.0176 |
| $\cos(\chi_{i})\sin(\chi_{2i})$ | 0.0272 | 0.0036 | 0.0058 | 0.0249 | 0.0106 | -0.0022 |
| $\cos(\phi_{i})\cdot \text{charge}_{i-1}$ | 0.0047 | -0.0222 | -0.0104 | 0.0176 | -0.0056 | -0.0111 |

| Attribute | $^{13}C^{\alpha}$ | $^{13}C^{\beta}$ | $^{13}C$ | $^{15}N$ | $^{1}H$ | $^{1}H^{\alpha}$ |
|---|---|---|---|---|---|---|
| electrostatic interaction shift | 0.0000 | 0.0000 | 0.0000 | -0.0436 | 0.0131 | 0.0122 |
| $\cos(\psi_{i})\sin(\psi_{i-1})$ | -0.0093 | 0.0141 | -0.0157 | -0.0027 | 0.0048 | -0.0204 |
| $\sin(\psi_{i}) \cdot \mathrm{charge}_{i+1}$ | 0.0031 | 0.0054 | -0.0037 | 0.0189 | 0.0050 | 0.0254 |
| $\sin(\psi_{i})\sin(\chi_{2i})$ | 0.0008 | -0.0168 | -0.0068 | -0.0216 | 0.0006 | -0.0145 |
| $\cos(\psi_{i})\sin(\chi_{2i})$ | 0.0060 | -0.0198 | -0.0094 | -0.0074 | -0.0004 | -0.0165 |
| $\cos(\phi_{i}) \cdot \mathrm{proline}_{i-1}$ | -0.0026 | 0.0015 | -0.0016 | -0.0060 | 0.0077 | 0.0339 |
| $\sin(\phi_{i}) \cdot \mathrm{charge}_{i-1}$ | -0.0057 | 0.0070 | -0.0073 | 0.0205 | 0.0087 | 0.0012 |
| $\sin(2\phi_{i-1})$ | -0.0096 | 0.0028 | -0.0029 | -0.0118 | 0.0021 | -0.0144 |
| $\sin(\psi_{i}) \cdot \mathrm{aromaticity}_{i+1}$ | 0.0020 | 0.0008 | -0.0028 | -0.0174 | 0.0092 | 0.0010 |
| $\cos(\phi_{i})\sin(\chi_{2i})$ | -0.0030 | 0.0018 | 0.0007 | 0.0012 | 0.0067 | 0.0007 |
| mean shift | 0.0000 | 0.0000 | 0.0000 | 0.0000 | 0.0000 | 0.0000 |
| methyl bond | 0.0000 | 0.0000 | 0.0000 | 0.0000 | 0.0000 | 0.0000 |
| contact sum | 0.0000 | 0.0000 | 0.0000 | 0.0000 | 0.0000 | 0.0000 |
| disulfide bond | 0.0000 | 0.0000 | 0.0000 | 0.0000 | 0.0000 | 0.0000 |

Table 5: A comparison of updated RNA chemical shift predictions using a newly-adopted FMs approach to a previously reported SVR model with analysis using the Automated-Plus approach [13]. The structure of the table is identical to that of Table 1 in [13], except that results for $^{15}$N nuclei, previously unconsidered, are reported for the FM approach. The database of shifts was also expanded with new RNA entries from the BMRB. RMSD values were calculated between the predicted and experimental shift values. $n$ denotes the number of shifts included in the calculation for the given category. [a]Ten-fold cross validation performed during model training for all atoms. [b]Canonical bases are the central base in a 5 base stretch in which all 5 base pairs have GC or AU base pairing and no other attributes such as being in a triplet, kissing interaction or pseudoknots are present. [c]Non-canonical bases are the same as canonical, but the first and/or fifth bases may be GU wobble base pairs, mismatched, unpaired (e.g. loops) or not-present (e.g. the 5′ or 3′ termini). [d]Other bases are all bases that are in neither the canonical nor non-canonical categories. [e]All denotes the average across all categories (b-d).

| Category | SVR | | FM | |
|---|---|---|---|---|
| | RMSD | $n$ | RMSD | $n$ |
| | | $^{1}$H | | |
| Cross-validated[a] | 0.13 | 18,774 | 0.11 | 38,099 |
| Canonical[b] | 0.06 | 3,020 | 0.09 | 7,422 |
| Non-canonical[c] | 0.07 | 2,903 | 0.09 | 6,120 |
| Other[d] | 0.11 | 12,851 | 0.11 | 24,557 |
| All[e] | 0.10 | 18,774 | 0.10 | 38,099 |
| | | $^{13}$C | | |
| Cross-validated | 0.83 | 9,642 | 0.84 | 17,956 |
| Canonical | 0.46 | 1,630 | 0.44 | 3,367 |
| Non-canonical | 0.47 | 1,526 | 0.49 | 2,786 |
| Other | 0.85 | 6,486 | 0.70 | 11,803 |
| All | 0.75 | 9,642 | 0.64 | 17,956 |
| | | $^{15}$N | | |
| Cross-validated | – | – | 1.32 | 3,460 |
| Canonical | – | – | 0.36 | 667 |
| Non-canonical | – | – | 0.33 | 536 |
| Other | – | – | 0.36 | 2,257 |
| All | – | – | 0.39 | 3,460 |

Table 6: NMRFx generates an output file, `analysis.txt`, after batch mode generation of multiple structures. The file lists the names of the involved atoms, the bound (upper or lower) that is violated, the number of structures with that violation, the mean value of that violation, the violation in the structure with the largest violation, and a text string where `+` indicates a structure with a violation.

<table>
<tr><th>Atom 1</th><th>Atom 2</th><th>$n_{\text{viol}}$</th><th>Bound</th><th>Mean</th><th>Max</th><th>Structures</th></tr>
<tr><td>1:31.H6</td><td>1:32.H6</td><td>3</td><td>5.00</td><td>0.13</td><td>0.28</td><td>|..++.....+|</td></tr>
<tr><td>1:29.H61</td><td>1:8.O4</td><td>6</td><td>2.00</td><td>0.22</td><td>0.28</td><td>|..++++..++|</td></tr>
<tr><td>1:10.H5′</td><td>1:10.H8</td><td>4</td><td>3.20</td><td>0.16</td><td>0.33</td><td>|...++++...|</td></tr>
<tr><td>1:10.P</td><td>1:18.P</td><td>9</td><td>13.00</td><td>0.26</td><td>0.37</td><td>|++.+++++++|</td></tr>
<tr><td>1:2.H8</td><td>1:3.H6</td><td>10</td><td>5.00</td><td>0.46</td><td>0.52</td><td>|++++++++++|</td></tr>
<tr><td>1:3.H3</td><td>1:34.N1</td><td>6</td><td>1.92</td><td>0.20</td><td>0.31</td><td>|+.+.++..++|</td></tr>
<tr><td>1:25.H3′</td><td>1:25.H8</td><td>6</td><td>3.50</td><td>0.47</td><td>0.82</td><td>|+++.+...++|</td></tr>
<tr><td>1:28.N3</td><td>1:9.H1</td><td>8</td><td>1.89</td><td>0.24</td><td>0.33</td><td>|+.+++++.++|</td></tr>
<tr><td>1:22.H6</td><td>1:23.H6</td><td>5</td><td>5.00</td><td>0.24</td><td>0.46</td><td>|+.+.+...++|</td></tr>
<tr><td>1:34.H8</td><td>1:35.H6</td><td>8</td><td>5.00</td><td>0.25</td><td>0.40</td><td>|++.++.++++|</td></tr>
<tr><td>1:27.P</td><td>1:4.P</td><td>4</td><td>11.00</td><td>0.21</td><td>−0.30</td><td>|+.....+++.|</td></tr>
<tr><td>1:10.H8</td><td>1:11.H8</td><td>8</td><td>5.00</td><td>0.44</td><td>0.62</td><td>|++.++++.++|</td></tr>
<tr><td>1:10.H5″</td><td>1:10.H8</td><td>7</td><td>3.80</td><td>0.18</td><td>0.28</td><td>|...+++++++|</td></tr>
</table>

### 5 Code Listings

**Code Listing 1:** Jython script (called `process.py` by default) used to process the NUS HNCACB dataset presented in Figure 1 of the main text. A script of this form (minus the comments which have been added for edification) can be generated indirectly using the processing accordion from the NMRFx GUI.

```
# Processing operations are exposed by NMRFx's `pyproc` library
from pyproc import *

# Use 10 processor cores.
# By default, half of the total number of cores are used.
procOpts(nprocess=10)
FID("/path/to/raw/nus/data")  # Load data
CREATE("/path/to/raw/nus/data/spec.nv")  # Create new NMRFx dataset
acqOrder("21")
acqarray(0, 0, 0)
fixdsp(True)
label("HN", "15N", "13C")
ref("", "", "")
skip(False, False, False)
acqmode("complex", "hyper", "hyper")

# === 1H dimension processing ===
DIM(1)
SUPPRESS()  # Water signal suppression
SB()  # Sine-bell apodization
ZF()  # Zero-filling
FT()  # Fourier Tranform
PHASE(ph0=-94.4, ph1=0.0)  # Phasing
EXTRACTP(start=10.75, end=5.9, mode="region")  # Region selection

# === Indirect matrix (15N & 13C) processing ===
DIM(2, 3)
# Specify NUS schedule
SCHEDULE("/path/to/raw/nus/data/nuslist.scd")
ZFMAT(zfy=1, zfz=1)  # Zero-filling
SB()
# `PHASE_ID` Fourier transforms the data, performs
# phase-correction, and then performs inverse Fourier
# transformation, returning the data to the time-domain
# for processing with GRINS.
# Each argument is a 2-tuple corresponding to the 15N
# and 13C dimensions, respectively.
# `negateImag` indicates whether the FID points should be
# transformed to their complex conjugates.
# `negatePairs` indicates whether every second FID point should
```

```
# be multiplied by -1.
PHASE_ID(
    ph0=[0.00, 0.00],
    ph1=[0.00, 0.00],
    negateImag=[False, False],
    negatePairs=[True, True],
)
# Perform NUS reconstruction with the GRINS algorithm
GRINS(noiseRatio=5.0)

# === 15N dimension processing ===
DIM(2)
SB()
ZF()
FT()

# === 13C dimension processing ===
DIM(3)
SB()
ZF()
FT()

# === Baseline correction (Whittaker smoothing) ===
DIM(1)
BCWHIT()
DIM(2)
BCWHIT()
DIM(3)
BCWHIT()

# === Run commands ===
run()
```

**Code Listing 2:** An example of a YAML file used to define the layout of spectrum charts in NMRFx. This layout is that used to generate the arrangement of NOESY, HMQC and TOCSY experiments in Figure 10.

```
layouts :
  - name : RNA-UMBC
    layout :

      # === Top-left panel ===
      - type : [noesy,tocsy]  # Datasets whose names contain
                              #   "noesy" or "tocsy" will be in
                              #   this chart
        loadpeaks : true      # If the dataset has an associated
                              #   peak list, load it
        row : 0               # Display in first row of the grid
        column : 0            # Display in first column of the grid
        rowspan : 2           # Chart spans 2 rows of the grid
        columnspan : 2        # Chart spans 2 columns of the grid
        x : [6.4, 8.5]        # X-axis spans 6.4 to 8.5 ppm
        y : [4.8, 6.2]        # Y-axis spans 4.8 to 6.2 ppm
        xsync : arom          # Synchronize x-axis of this chart
                              #   with other axes labelled "arom"
        ysync : h1py          # Synchronize y-axis of this chart
                              #   with other axes labelled "h1py"

      # === Bottom-left panel ===
      - type : [hmqc]
        loadpeaks : true
        row : 2
        column : 0
        rowspan : 1
        columnspan : 2
        x : [6.4, 8.5]
        y : [133.5, 157]
        xsync: arom

      # === Top-right panel ===
      - type : [noesy,tocsy]
        loadpeaks : true
        row : 0
        column : 2
        rowspan : 2
        columnspan : 1
        x : [4.8, 6.2]
        y : [4.8, 6.2]
        xsync : h1p
        ysync: h1py

```

```
      # === Bottom-right panel ===
      - type : [hmqc]
        loadpeaks : true
        row : 2
        column : 2
        rowspan : 1
        columnspan : 1
        x : [4.8, 6.2]
        y : [88, 106.5]
        xsync : h1p
```

**Code Listing 3:** The YAML file used to calculate the RNA structure. The file specifies the molecular topology, the files used for restraints, and the parameters used in the torsion angle dynamics and optimization.

```yaml
molecule :
  entities :
      - sequence : GGUUGAGUGGAACUGUGAAGUUCGGAACACUCAACC
        ptype : RNA
        chain : A

tree:

distances :
    - file : slcm-hb
      type : cyana
    - file : slcm-pp
      type : cyana
    - file : slcm-noe
      type : cyana

angles :
    - file : all.aco
      type : cyana

rdcs :
    - file : slca_rdcs_for_analyst.dat
      type : nmrfx

anneal:
    dynOptions :
        steps : 15000
        highTemp : 5000.0
        dfreeSteps : 10000
    force :
        tors : 0.1
        dih : 10
        irp : -0.2
        rdc : 0.5
    stage_low:
        force :
            repel : 4.0
```
